# Immunophenotyping of T cells Combined with Vβ antibodies identifies long lasting CMV related T cell Expansions with a consistent TIGIT and PD-1 phenotype

**DOI:** 10.1101/2024.08.30.610465

**Authors:** Monique M. van Ostaijen-ten Dam, Marco W. Schilham, Arjan C. Lankester, Janine E. Melsen

**Affiliations:** Laboratory for Pediatric Immunology, Willem-Alexander Children’s Hospital, Leiden University Medical Center, Leiden, The Netherlands; Department of Immunology, Leiden University Medical Center, Leiden, The Netherlands

**Author notes:** **Correspondence** Janine Melsen Albinusdreef 2, 2333 ZA The Netherlands.

## Abstract

Simultaneous analysis of T cell receptor repertoire and T cell phenotype is of high relevance to gain a deeper understanding on the T cell response in terms of differentiation, kinetics and persistence. Also it is currently unknown how the repertoire of immune checkpoints on T cells evolves over time. To fully capture the heterogenous response, TCR repertoire and T cell phenotype analysis is preferably performed at the single-cell level, but this is costly and analytically challenging. Therefore, we developed a flow cytometry based method that allows for simultaneous characterization of the T cell TCR Vβ and the phenotype. We generated a 9-tube 24-color panel with antibodies against maturation and activation markers, and 24 TCR Vβ chains. Peripheral blood mononuclear cells from 5 healthy controls were analyzed, revealing the presence of oligoclonal expansions. Within the CD27+CD28+ memory T cell population, oligoclonal expansions were characterized by a T regulatory, or KLRG1+CCR7-CD27low phenotype. Within the late CD27- and/or CD28-memory T cells, the populations with the lowest TCR Vβ diversity expressed CX3CR1. As a proof of principle, we demonstrate the dynamics of circulating oligoclonal T cell expansions that emerged following hematopoietic stem cell transplantation in the presence of CMV in 2 patients. We provide evidence that oligoclonal T cells transferred from donor to recipient can be traced up to at least 2.5 years while maintaining its phenotype. Mostly TIGIT and PD-1 were critical in defining T cell expansions, while CD45RA, CD57, CD56 and NKG2A were variably expressed. Overall, we developed a phenotype based method to trace T cell populations, that is widely applicable to different settings in which the T cell response needs to be monitored.

## Introduction

T cells are adaptive lymphocytes which are critical for the defense against a variety of pathogens. Each T cell is endowed with a T cell receptor (TCR), which is in more than 90% of the T cells composed of an alpha (α) and a beta (β) chain. During thymic development, the chains of the TCR are genetically rearranged by random recombination of the Variable (V), Diversity (D) and Joining (J) regions, and insertion of nucleotides.^1^ This process ensures a highly diverse repertoire of unique TCRs to combat a broad spectrum of pathogens. Three hypervariable complementarity determining regions (CDRs) of the rearranged TCR chains form the antigen binding site, of which the most variable CDR3 determines the antigen specificity of the TCR.^2^ Following recognition of a foreign antigen in the context of major histocompatibility complex (MHC), T cells respond and expand, a process known as clonal selection.^3^

Identification and monitoring of protective or pathogenic T cell clones could be relevant in different settings, such as vaccination, viral infection, autoimmune disease or transplantation. Multiple reports demonstrated that distinct antigenic stimuli imprint distinct phenotypical profiles in the reactive T cells, suggesting that memory formation is a heterogenous response.^4–6^ For instance, Dengue, CMV-, EBV- and Flu-specific CD8^+^ T cells have been associated with different maturation stages, as has been shown by combined tetramer and maturation marker staining using CyTOF, and by cellular indexing of transcriptomes and epitopes (CITE-seq) combined with TCR receptor sequencing.^7–9^ CMV-specific CD8 memory T cells are characterized by loss of CD27 and CD28, while expressing CD57, CX3CR1 and granzyme B. In contrast, EBV-specific CD8 memory T cells still express CD27 and CD28. In addition to these markers, chronic viral activation has been associated with upregulation of a multitude of inhibitory receptors on T cells, such as PD-1 and TIGIT.^10,11^ How these maturation, activation and exhaustion phenotypes develop longitudinally on each clonal T cell subset, from antigen exposure to clearance or chronic persistence, remains poorly understood.

To unravel the heterogenous response, TCR repertoire and T cell phenotype analysis is preferably performed at the single-cell level. The recent introduction of single-cell RNA sequencing, combined with TCR sequencing and protein expression, allows detailed exploration of the T cell compartment. However, since this approach is costly and analytically difficult, application as a routine or diagnostic tool is unfeasible. In this study, we aimed to explore a robust cytometry based approach. We generated a 9-tube 24-color panel with antibodies against maturation and activation markers. The TCRβ repertoire was assessed by antibodies against TCR Vβ chains. As a proof of principle, we demonstrate the dynamics of T cell expansions following hematopoietic stem cell transplantation in the context of CMV.

## Materials and Methods

### Ethics statement

Blood samples from two hematopoietic stem cell transplant (HSCT) recipients, its two healthy donors and three healthy controls (Supplementary Table 1) were drawn after informed consent was provided. HSCT recipient 1 received bone marrow from a matched family donor as treatment for β-thalassemia. HSCT recipient 2 received bone marrow from an matched sibling donor as treatment for Shwachman-Diamond syndrome. Both patients received reduced intensity conditioning prior to HSCT (Fludarabine, Busulfex and ATG). Approval for the study was given by the Institutional Review Board (B.17.001). Peripheral blood mononuclear cells (PBMCs) were isolated by Ficoll density gradient and analyzed after cryopreservation.

### Spectral Cytometry

PBMC were stained with fluorochrome-conjugated antibodies for 30 minutes at room temperature (RT) in PBS supplemented with 0.5% BSA, 2mM EDTA, 0.02% NaN3 and Brilliant Stain Buffer plus (Becton Dickinson (BD), Franklin Lakes, NJ, USA). The panel consisted of 8 tubes, each including three distinct FITC and/or PE conjugated Vβ antibodies (Beckman Coulter, Brea, CA, USA) and a backbone panel of surface markers. The ninth tube combined CD45RO (FITC) and TIM-3 (PE) with the backbone panel. All the antibodies used are listed in Supplementary table 2. 7AAD was added prior to acquisition to exclude dead cells. Data was acquired on a 3L Aurora spectral cytometer (Cytek Biosciences, Fremont, CA, USA) using SpectroFlo software (v2.0, Cytek).

### Data analysis

The OMIQ platform (www.omiq.ai, Dotmatics, Boston, MA, USA) was used for the post-acquisition analysis of cytometry data (Figure 1). As part of the preprocessing, data was compensated and arcsinh transformed using individual cofactors per parameter^12^. Living single CD3+ T cells were gated and flowAI was applied to remove unwanted anomalies, based on changes in flowrate and outlier events.^13^ Some additional outlier events (<1% of the T cells) based on extremely high expression of CD127, not detected by FlowAI, were removed by manual gating. Next, γδ T cells were excluded and CD4+ and CD8+ T cells were gated and subsampled to include a similar amount of cells per tube. Based on FITC and PE expression within the CD4 and CD8 T cells, four different populations were gated: three Vβ+ and one Vβ-population. Subsampling among the 8 tubes within each sample was performed to ensure an equal number of total αβ T cells per tube within one sample. The naïve and memory populations were defined based on either a combination of CCR7, CD45RA and CD95 only or with addition of CD27 and CD28. The early CD27^+^CD28^+^ memory and late CD27^-^ and/or CD28^-^ memory T cells were selected for opt-SNE^14^ based on all markers except CD3, TCRgd, CD16, CD4, CD45RO, TIM-3 and Vβs. Clustering was performed by ClusterX^15^ (distance cutoff 3) based on the two Opt-SNE coordinates. The CD8 late memory T cells of HSCT1 were downsampled to 1 million cells to enable clustering. Heatmaps and stacked backgraphs were generated in R (v4.4.1, R Foundation for Statistical Computing, Vienna, Austria), using the pheatmap^16^ and ggplot2 package^17^, respectively. The Inverse Simpson index^18^ was calculated using the package Abdiv^19^. Other graphs were created in Graphpad Prism software (v10.2.3, GraphPad Software, Boston, MA, USA).

**Figure 1.**
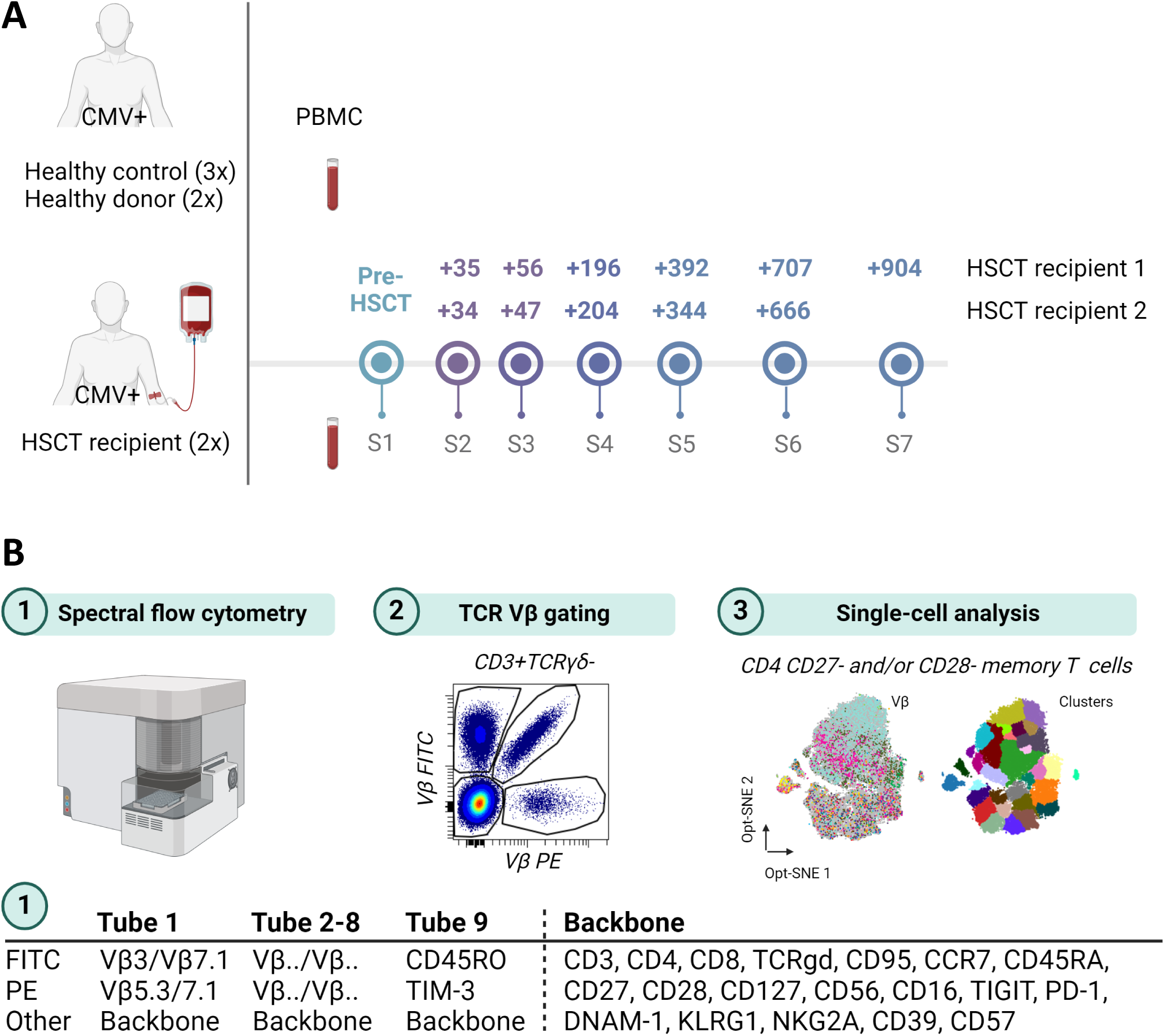
Overview analysis pipeline. **A.** Peripheral blood mononuclear cells (PBMC) from three CMV seropositive healthy controls, and two CMV seropositive healthy donors were included. Longitudinal PBMC samples from two CMV seropositive hematopoietic stem cell transplant (HSCT) recipients were included. **B.** A 9-tube spectral flow cytometry panel was developed including Vβ antibodies (3 per tube, except for tube 9) and a backbone panel of markers. After preprocessing of the data the individual Vβ populations were labeled and single-cell analysis (dimensionality reduction and clustering) was applied on different populations of CD4 and CD8 T cells. Created with BioRender.com

## Results

### Loss of CD27 and/or CD28 characterizes memory T cell population with reduced TCR Vβ diversity

To study the phenotype of the T cell compartment combined with the TCR Vβ repertoire, a 24-color spectral cytometry panel was developed. First, 5 healthy blood samples from CMV seropositive healthy controls (HC) and healthy donors (HD) ranging in age from 2-47 years old were analysed (Figure 1A). The analysis pipeline included gating of the TCR Vβ populations, dimensionality reduction by opt-SNE and density based clustering by ClusterX on different population of T cells (Figure 1B). Within the total CD4 and CD8 compartment we identified the naïve (CD95-CCR7+CD45RA+), central memory (CM, CD95+CCR7+CD45RA-), effector memory (EM, CD95+CCR7-CD45RA-) and effector memory RA+ (EMRA, CD95+CCR7-CD45RA+) population using the conventional definition (Figure 2A).^20^ Since we observed previously within clonal T cell expansions gradients of CD45RA, we hypothesized that use of CD27 and CD28 to label memory populations, would improve identification of clonal populations.^21^ Therefore, as an alternative definition, we also categorized the memory populations into early memory (CD27+CD28) and late memory (CD27- and/or CD28-) T cells.

**Figure 2.**
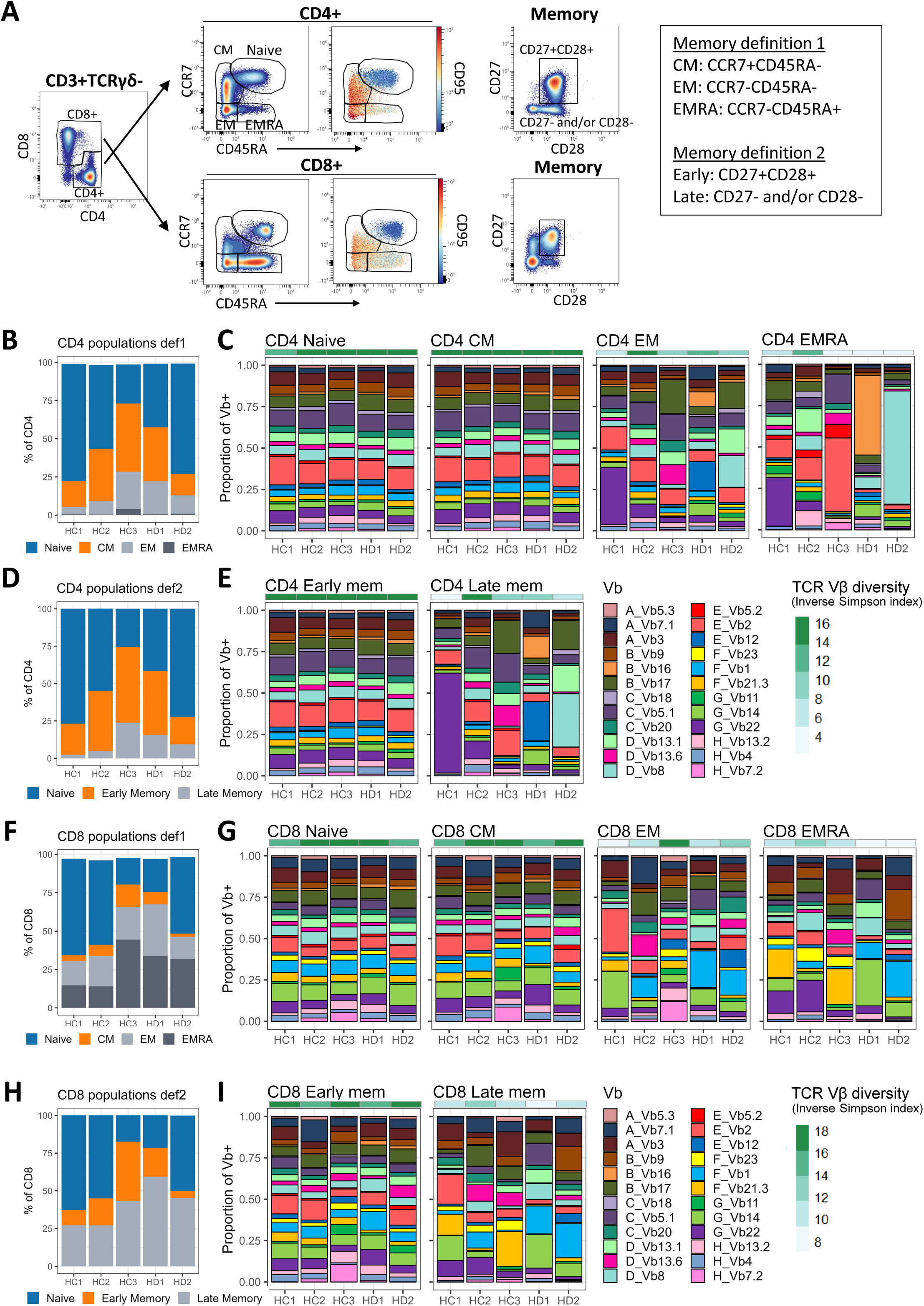
TCR Vβ distribution in healthy donors per T cell population using two definitions. **A.** The CD4 and CD8 T cells were gated, followed by gating of the naïve T cells defined as CD95+CCR7+CD45RA+. The remaining CD4 or CD8 T cells were either categorized as central memory (CM), effector memory (EM) and effector memory RA+ (EMRA) using CCR7 and CD45RA (definition 1), or as early and late memory based on CD27 and CD28 (definition 2). **B.** The distribution of the CD4 naïve and memory populations (definition 1) in the individual control samples. **C.** The TCR Vβ distribution per individual donor, per CD4 naïve and memory compartment. The colorbar at the top represents the inverse Simpson index (TCR Vβ diversity). **D.** The distribution of the CD4 naïve and memory populations (definition 2) in the individual control samples. **E.** The TCR Vβ distribution per individual donor, per CD4 memory compartment (definition 2). **F.** The distribution of the CD8 naïve and memory populations (definition 1) in the individual control samples. **G.** The TCR Vβ distribution per individual donor, per CD8 naïve and memory compartment. The colorbar at the top represents the inverse Simpson index (TCR Vβ diversity). **H.** The distribution of the CD8 naïve and memory populations (definition 2) in the individual control samples. **I.** The TCR Vβ distribution per individual donor, per CD8 memory compartment (definition 2).

The TCR Vβ distribution of the total CD4 T cells was similar to the reference values (based on at least 46 donors) as reported by the antibody kit (Figure S1A). We confirmed that the 24 TCR Vβ antibodies cover approximately 70% of the TCR Vβ repertoire (Figure S1). Within the CD4 compartment we detected variable frequencies of the different memory populations, with the EM and EMRA population being most prevalent in HC3 (24.4% and 4.1% respectively, Figure 2B). When zooming on the different subsets, we detected a similar diverse Vβ distribution between total, naïve and CM CD4 T cells (Figure S1A), as also reflected by the inverse Simpson index (diversity score, Figure 2C). The EM and EMRA population of all healthy individuals, except HC2, were less diverse. When using the alternative memory definition based on CD27 and CD28 it became clear that the late memory CD4 T cells reflected the altered Vβ distribution that was observed in the EM and/or EMRA population (Figure 2D,E). For the CD8 T cells, HC3 also had the highest percentage of EMRA CD8 T cells (44.4%, Figure 2F). The distribution of the total, naïve and CM CD8 compartment was highly similar to the reference values from the kit (Figure 2G, S1C). Like for the CD4 T cells, the CD8 TCR Vβ diversity was reduced in the EM, EMRA and late memory CD8 compartment. Since the late memory population in general captured the oligoclonal expansions that were present in both the EM and EMRA compartment, we decided to continue our analysis using the early/late memory definitions.

### In the early memory CD4 T cell compartment a few TCR Vβ dominant clusters can be detected mostly with a KLRG1+CCR7-CD27low or Treg phenotype

As a next step, we zoomed in on the early memory T cells to unravel the TCR Vβ repertoire and its association with its phenotype. An Opt-SNE of every individual donor was generated and the cells were clustered (Figure 3A, S2). The diversity of the TCR Vβs (inverse Simpson index) and the Euclidean distance to the naïve CD4 T cells was calculated based on the percentage of the Vβs in every cluster. A cluster with oligoclonal expansions (and thus dominant Vβs) was defined as >0.2% of all Vβ+ cells in the cluster, TCR Vβ diversity <11 and distance to naïve CD4 T cells >1.4. In agreement with the naïve-like TCR Vβ diversity, only a few Vβ dominant clusters with an early memory phenotype were detected (Figure 3B). The clusters with the lowest TCR Vβ diversity expressed KLRG1 and did not express CCR7 (C01_HD2, C14_HC2, C23_HC3, C49_HD1, C37_HC3, C12_HC3, C40_HD2, C44_HD2, Figure 3C, Figure S2). Although these clusters were positive for CD27 and CD28, the median signal of CD27 was lower compared to the CD4 Naïve T cells, suggesting that they are starting to loose CD27 (Figure 3C). In addition, two clusters with a reduced TCR Vβ diversity had a regulatory T cell phenotype (CD127low and CD39+, CD24_HD2, CD23_HD2).

**Figure 3.**
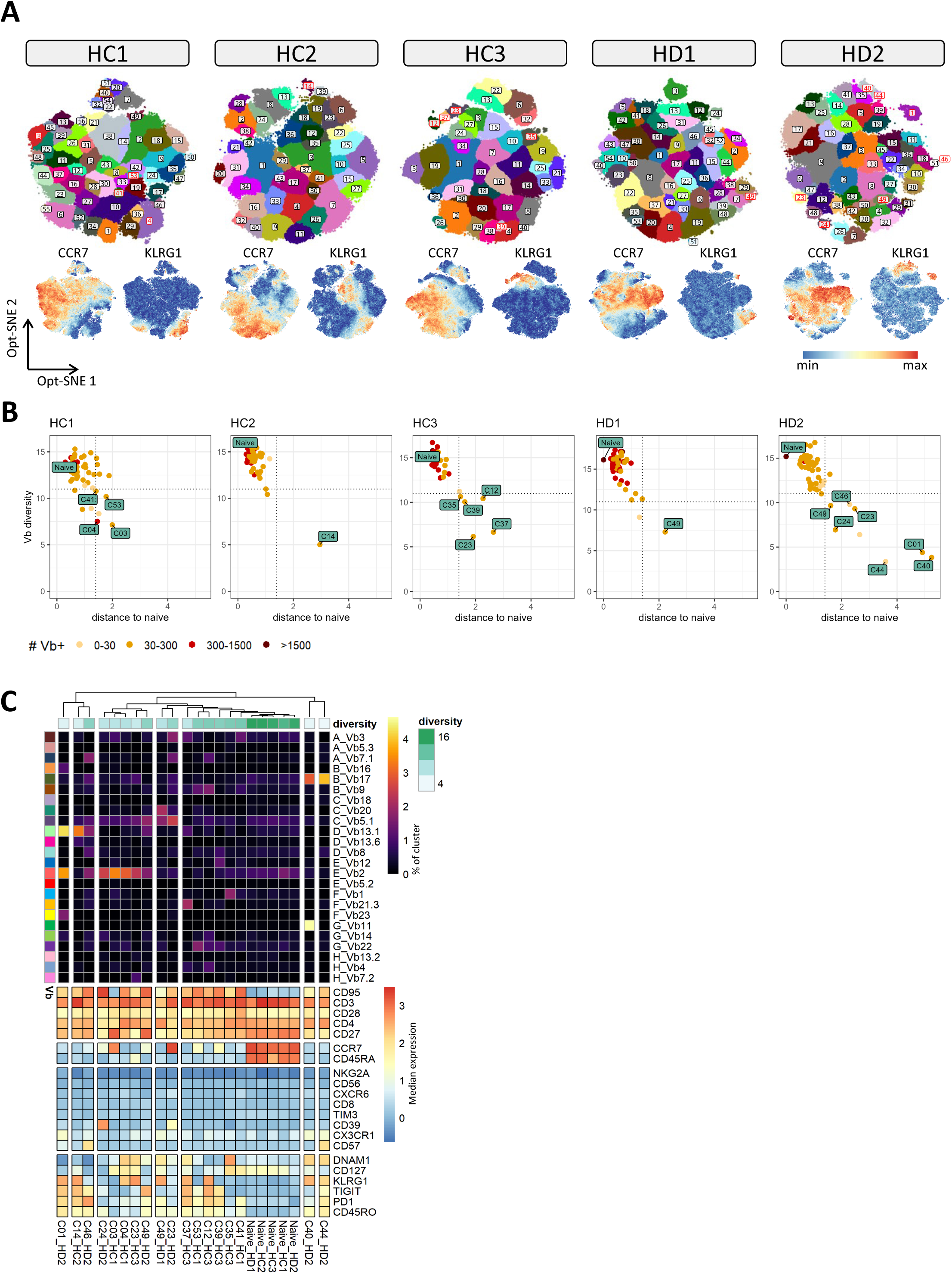
Oligoclonal expansions in the early memory CD4 compartment have a KLRG1+ or Treg phenotype. **A.** The early memory CD4 T cells of each healthy control (HC) and donor (HD) were embedded in an Opt-SNE and clustered by ClusterX. The clusters that were defined as oligoclonal expansion in B, are labeled in red. **B.** For every cluster the TCR Vβ diversity (inverse Simpson index) and the Euclidean distance to the naïve CD4 T cells were calculated based on the frequencies of the individual Vβs per cluster. A cluster with oligoclonal expansions was defined as Vβ diversity <11, distance >1.4 and total number of Vβ+ cells >0,2% of all Vβ+ cells. The naïve cluster and the clusters that meet these criteria are labeled. **C.** The frequency of the Vβs (% of total cells in cluster) and the phenotype per cluster (median intensity), as labeled in B. As a reference the naïve CD4 T cells are included. The diversity represents the inverse Simpson index.

### All CX3CR1+ CD4 late memory T cells are defined as oligoclonal expansion and cluster based on TIGIT and PD-1

In contrast to the CD4 early memory T cells, within the CD4 late memory T cells the vast majority of clusters (except in HC2) were defined as oligoclonal expansion (Figure 4A,B). Moreover, in most of the clusters only one or two dominant TCR Vβs were detected (Figure 4A,C). In HC1, the TCR Vβ22 enriched clusters even dominated the whole compartment since they represented 53% of all late memory CD4 T cells (Figure 1E, 4A). Overall, the TCR Vβ dominant clusters were characterized by expression of CX3CR1, a chemokine receptor reported on CMV-specific CD8 T cells (Figure 4A, 4C, S3). The clusters in which the same Vβ was dominant, heterogeneously expressed CD57, CD8 and/or CD45RA (e.g. TCR Vβ22 and Vβ12 enriched clusters, Figure 4C). CD56 was observed on some clusters derived from HC3 and HD1, highlighting the importance of including CD3+CD56+ T cells in the analysis. NKG2A was negative on all CD4 memory T cells.

**Figure 4.**
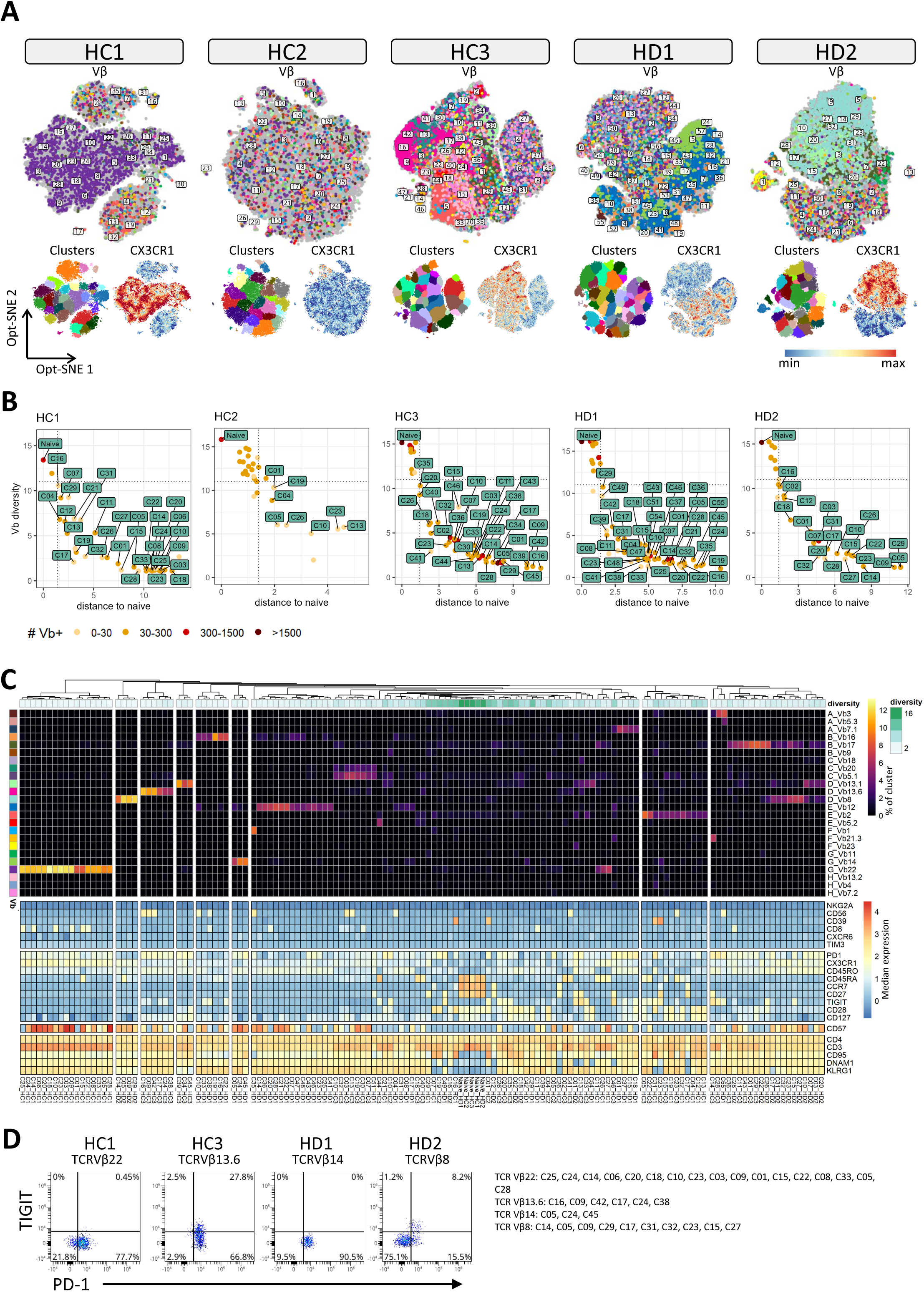
Clusters of the late memory CD4 compartment with only 1 or 2 dominant Vβs express CX3CR1. **A.** The late memory CD4 T cells of each healthy control (HC) and donor (HD) were embedded in an Opt-SNE and clustered by ClusterX. Each cell is colored by its TCR Vβ expression, cluster assignment, or CX3CR1 expression. **B.** For every cluster the TCR Vβ diversity (inverse Simpson index) and the Euclidean distance to the naïve CD4 T cells were calculated based on the frequencies of the individual Vβs per cluster. A cluster with oligoclonal expansions was defined as Vβ diversity <11, distance >1.4 and total number of Vβ+ cells >0,2% of all Vβ+ cells. The naïve cluster and the clusters that meet these criteria are labeled. **C.** The frequency of the Vβs (% of total cells in cluster) and the phenotype per cluster (median intensity, as labeled in B. As a reference the naïve CD4 T cells are included. **D.** The TCR Vβ+ cells from the respective TCR Vβ dominant clusters were selected to study the TIGIT and PD-1 expression.

Previously, we observed in a HSCT patient with HPV-related clonal T cells expansions, that each clonal expansion had a homogenous phenotype of TIGIT and PD-1.^22^ In line with this, the TCR Vβ22 (HC1), Vβ13.6 (HC3) and Vβ14 (HD1) dominant clusters were nearly all TIGIT negative and PD-1 positive (Figure 4D). A minor fraction of TCR Vβ13.6+ cells from the respective clusters from HC3 also expressed TIGIT. The Vβ8 dominant clusters from HD2 were mostly TIGIT-PD-1-, but also some TIGIT+PD-1+, TIGIT-PD-1+ cells were observed (Figure 4D, S3). Overall, the vast majority of cells within a cluster defined as oligoclonal expansion is characterized by CX3CR1 expression, a homogenous TIGIT and PD-1 phenotype, and variable levels of CD45RA, CD8, CD57 and CD56.

### KLRG1 is abundantly expressed by CD8 early memory T cell expansions and defines oligoclonal expansions

Although the overall TCR Vβ diversity of CD8 early memory T cells was nearly identical to CD8 Naïve T cells, many clusters were identified in which the TCR Vβ distribution was altered. Compared to the CD4 early memory T cells KLRG1 was much more abundant (Figure 3A, 5A). Indeed, the vast majority of clusters defined as an oligoclonal expansion expressed KLRG1 (Figure 5B, C). Amongst the few TCR Vβ dominant clusters that did not express KLRG1, clusters were included that were positive for CD4 and CD39, potentially reflecting Tregs. While being absent on CD4 early memory T cells, NKG2A and CD56 were detected on some CD8 Vβ dominant clusters (Figure 5C, S4). With the exception of C35-HD1, PD-1 and TIGIT were again consistently expressed in clusters with the same dominant TCRVβ (Figure 5C).

**Figure 5.**
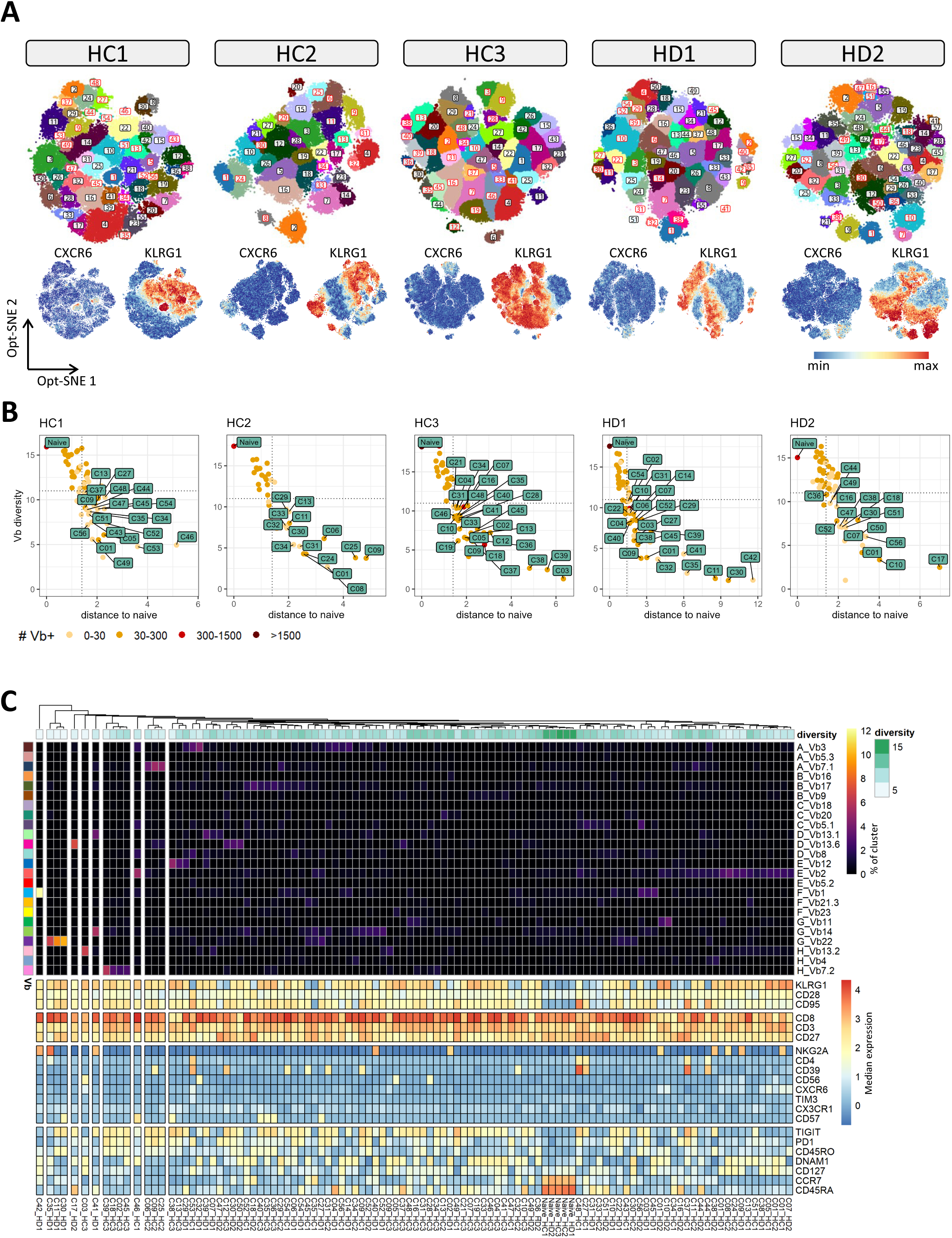
KLRG1 defines oligoclonal T cells expansions in early memory CD8 T cell compartment. **A.** The early memory CD8 T cells of each healthy control (HC) and donor (HD) were embedded in an Opt-SNE and clustered by ClusterX. The clusters that were defined as oligoclonal expansion in B, are labeled in red. **B.** For every cluster the TCR Vβ diversity (inverse Simpson index) and the Euclidean distance to the naïve CD8 T cells were calculated based on the frequencies of the individual Vβs per cluster. A cluster with oligoclonal expansions was defined as Vβ diversity <11, distance >1.4 and total number of Vβ+ cells >0.02% of all Vβ+ cells. The naïve cluster and the clusters that meet these criteria are labeled. **C.** The frequency of the TCR Vβs (% of total cells in cluster) and the phenotype (median intensity) per cluster, as labeled in B. As a reference the naïve CD8 T cells are included.

Clusters that expressed low levels of CXCR6 in the presence or absence of NKG2A were all enriched for either Vβ13.2 (C03_HC3), Vβ11 (C01_HD2, C10_HD2), or Vβ2 (C01_HD1, C08_HC2, C24_HC2, C09_HD1, C38_HD1, C05_HC1, C01_HC2, C01_HC1, C07_HD2, Figure 5A, 5C, S4). Vβ2 and Vβ13 are preferentially used by mucosal-associated invariant T (MAIT) cells, suggesting that these clusters represent the CXCR6+ circulating MAIT cells.^23–25^

### TCR Vβ dominant clusters in the CD8 late memory T cell compartment express CX3CR1 and have a heterogenous CD45RA, CD57, CD56 and NKG2A phenotype

For the CD8 late memory we observed a similar pattern as for the CD4 late memory T cells. All the CX3CR1 positive cells were defined as an oligoclonal expansion (Figure 6A, B). Again, in most of the clusters only one or two dominant TCR Vβs were detected (Figure 6C). Although CD8 T cells were described to first loose CD28 followed by CD27 during differentiation, we also detected clusters that were CD27-CD28+ (e.g. C12_HC3, C47_HC3, C13_HD1, C60_HC3, C02_HC3, C53_HC3, C05_HC2, Figure 6C, S5). This suggests that that not every T cell follows the same path of differentiation. PD-1 was homogenously expressed, but a bimodal expression of TIGIT was observed between clusters with the same dominant Vβ. To further analyze this phenotypic diversity, we selected the TCR Vβ1+ cells from the HD1 and HD2 clusters in which Vβ1 was dominant, and gated the TIGIT+ and TIGIT-cells. Interestingly, in HD1, CD45RA was specifically expressed on a fraction of the TIGIT+ cells, while CD56 was only expressed by a fraction of the TIGIT-cells, suggesting the presence of different clones (Figure 6D). In HD2, the phenotype of TIGIT+ and TIGIT-cells was reminiscent, thus it remains to be studied whether these clusters represent one clone with a variable TIGIT expression, or multiple clones. In addition to CD45RA and CD56, the markers NKG2A and CD57 were variably expressed between clusters with the same dominant TCR Vβ (Figure 6C).

**Figure 6.**
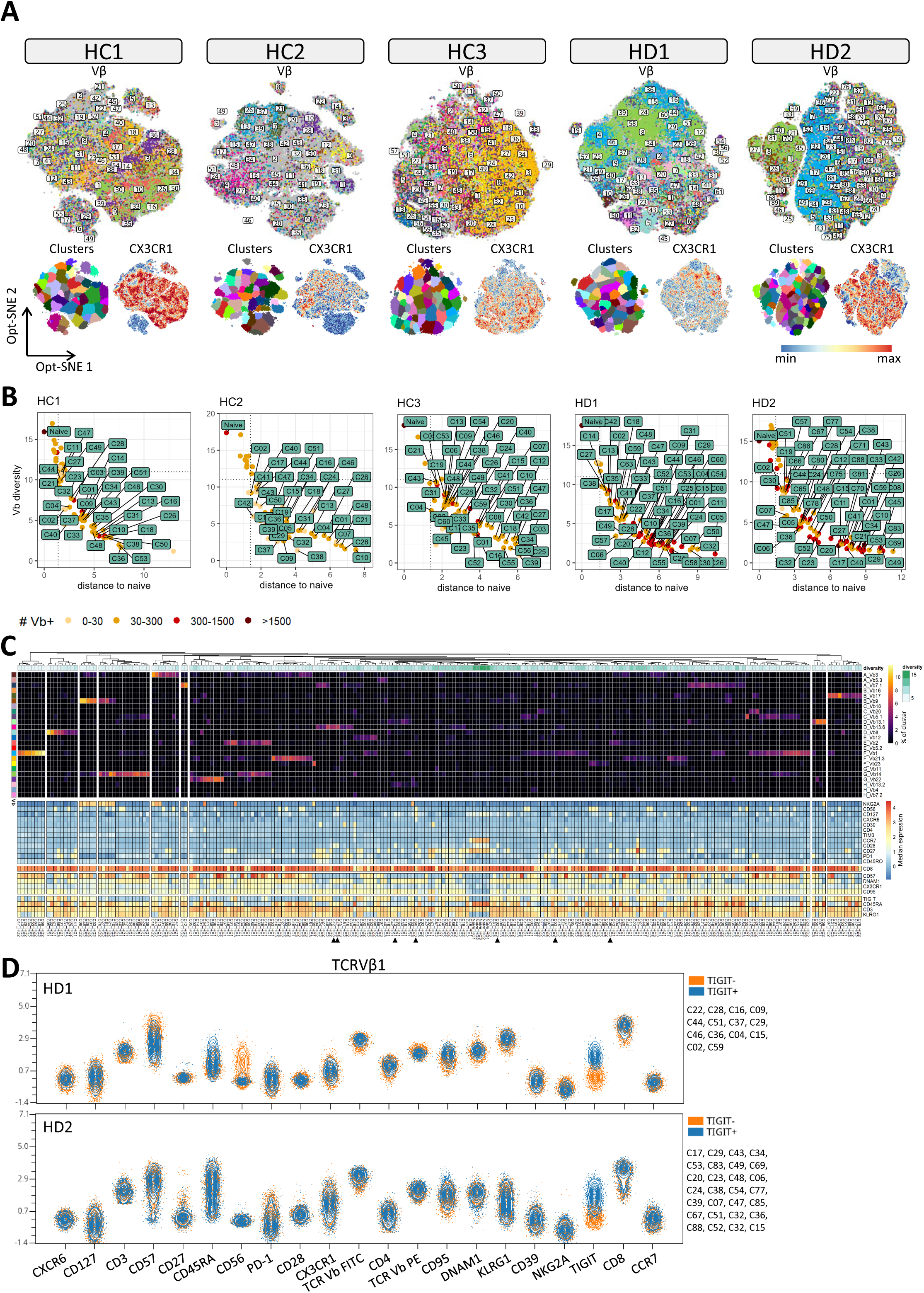
Oligoclonal expansions in the late memory CD8 T cell compartment express CX3CR1. **A.** The late memory CD8 T cells of each healthy control (HC) and donor (HD) were embedded in an Opt-SNE and clustered by ClusterX. Each cell is colored by its TCR Vβ expression, cluster assignment, or CX3CR1 expression **B.** For every cluster the TCR Vβ diversity (inverse Simpson index) and the Euclidean distance to the naïve CD8 T cells were calculated based on the frequencies of the individual Vβs per cluster. A cluster with oligoclonal expansions was defined as Vβ diversity <11, distance >1.4 and total number of Vβ+ cells >0.2% of all Vβ+ cells. The naïve cluster and the clusters that meet these criteria are labeled. **C.** The frequency of the Vβs (% of total cells in cluster) and the phenotype (median intensity) per cluster, as labeled in B. As a reference the naïve CD8 T cells are included. The arrowheads point towards the CD27-CD28+ clusters **D.** The clusters from HD1 and HD2 in which TCR Vβ1 was dominant were pooled per donor and separated into a TIGIT+ and TIGIT-population. The different markers are visualized per population.

### CMV related CD8 T cell expansions can be traced based on their phenotype and TCR Vβ until at least 1,5 years after hematopoietic stem cell transplantation

The CX3CR1 positive CD4 and CD8 late memory T cell expansions as observed in the healthy controls and donors, are most likely CMV-driven. To study the kinetics and phenotype of CMV related T cell expansions we longitudinally profiled PBMC samples from two HSCT recipients that were CMV seropositive before HSCT, and were transplanted with a CMV seropositive donor. In HSCT recipient 1 (hereafter referred to as HSCT1) at day 34 post transplantation a peak in the CMV load in the plasma was detected. At this timepoint also an expansion of the CD4 late memory (209 cells/µl) and CD8 late memory T cells (147 cells/µl) was observed (Figure 7A,B). The same approach was applied to identify clusters with oligoclonal expansions as was used in the healthy controls/donors. Data of the longitudinal samples were initially combined for the clustering. Although in the CD4 early memory T cell compartment only 2 clusters were detected with TCR Vβ expansions, in the CD4 late memory, CD8 early memory and CD8 late memory, the vast majority of clusters were defined as oligoclonal expansion (Figure 7C).

**Figure 7.**
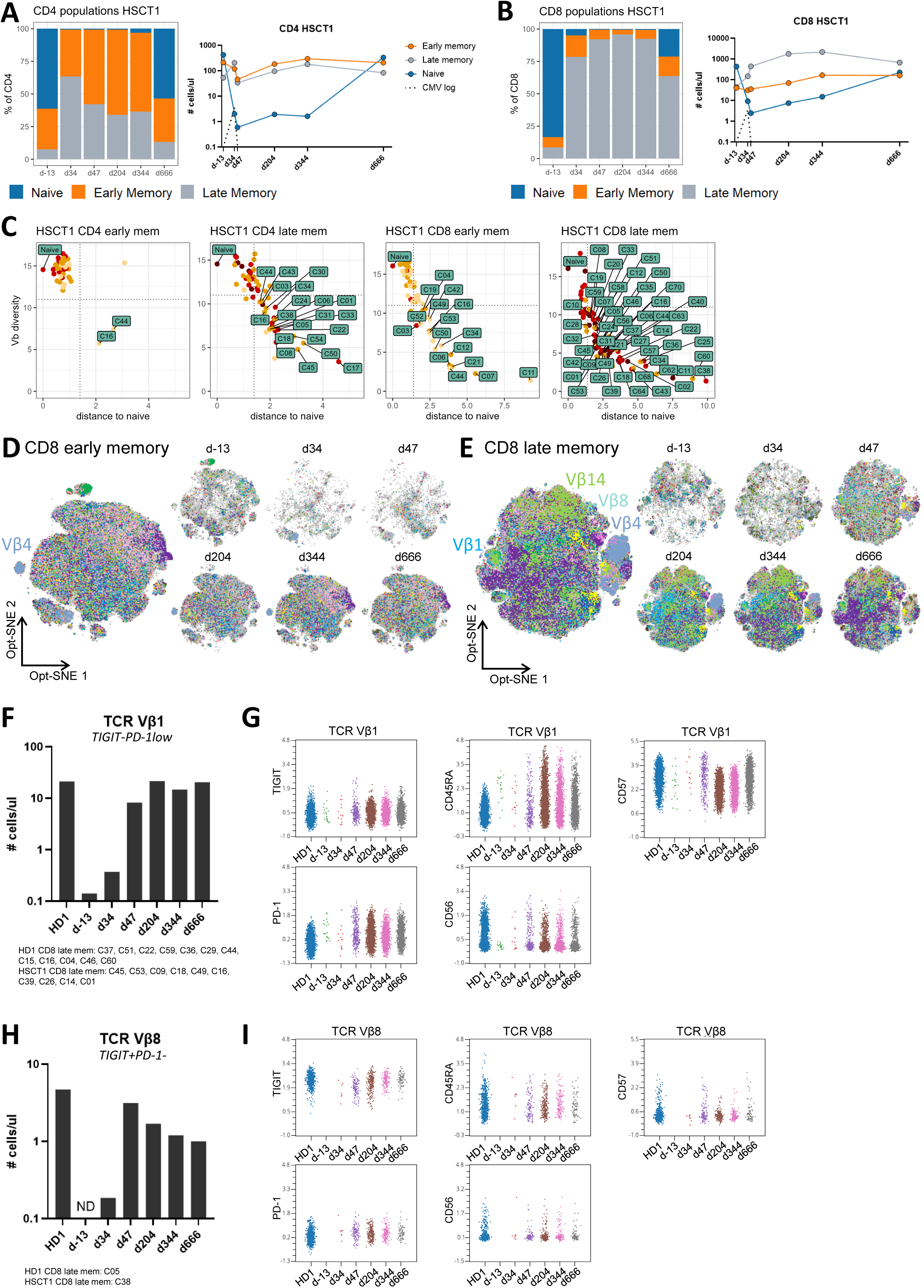
Oligoclonal T cell expansions can be traced until at least 1.5 years after HSCT in patient 1. **A.** Frequency and absolute numbers of CD4 and **B.** CD8 populations in PBMC samples before and after hematopoietic stem cell transplantation (HSCT). The dotted line represent the CMV load. **C.** The samples from HSCT1 were merged and an Opt-SNE and Opt-SNE based clustering was performed on the distinct CD4 and CD8 memory populations. For every cluster the TCR Vβ diversity (inverse Simpson index) and the Euclidean distance to the naïve T cells from before the HSCT were calculated based on the frequencies of the individual Vβs per cluster. A cluster with oligoclonal expansions was defined as Vβ diversity <11, distance >1.4 and total number of Vβ+ cells >0.2% of all Vβ+ cells. The naïve cluster and the clusters that meet these criteria are labeled. **D.** Opt-SNE visualization of the CD8 early and **E.** CD8 late memory T cells. Each cell is colored by its TCR Vβ expression. **F.** From the CD8 late memory clusters with a dominance of CD8 TCR Vβ1 and highest frequency at d47, the TCR Vβ1+ cells were selected and quantified. **G.** A consistent TIGIT and PD-1 (TIGIT-PD-1low) was observed over time, but a variable CD45RA and CD56 expression. **I.** From the CD8 late memory clusters with a dominance of CD8 TCR Vβ8 and highest frequency at d34/d47, the TCR Vβ8+ cells were selected and quantified. ND=not detected **J.** A consistent TIGIT and PD-1 (TIGIT+PD-1-) was observed over time, but a variable CD45RA and CD56 expression.

The T cell expansions around d34 and 47 are most likely caused by CMV reactivation. Since at this timepoint no new naïve T cells were formed in the patient, these expansions were mostly likely derived from donor memory T cells in the graft or from pre-existing memory T cells in de patient. In the CD4 T cells, only four clusters had its highest frequency at day 34 post HSCT (Figure S6A).These CD4 late memory had a phenotype, rarely seen in healthy individuals or before HSCT since they were negative for KLRG1, highly expressed PD-1 and expressed TIM-3 (Figure S6A,B). This phenotype was specific for the majority of all CD4 late memory T cells in the d34 sample, and not restricted to one TCR Vβ (Figure S6B, S6C, S7). Therefore, the cytokine rich environment shortly after HSCT in combination with the CMV reactivation potentially induces a specific phenotype on the CD4 late memory T cells.

For the CD8 T cells, TCR Vβ1 and TCR Vβ8 dominant clusters were detected in in both patient and the donor with a resembling phenotype (Figure S8, 7D,7E). In HSCT1, Vβ1+ cells were abundant from d47 onwards in the late compartment (Figure S8, 7F). The phenotype of these cells was similar between the donor and the recipient over time, being TIGIT-PD-1low, with a variable expression of CD45RA, CD56 and CD57 (Figure S7, S8, 7G). The TCR Vβ8+ cells, that were also abundant from d47 onwards, were mostly TIGIT+PD-1- and CD57-, with a variable expression of CD45RA and CD56 (Figure 7H, I). Overall, CMV related T cell expansions with a consistent phenotype, can be traced for a minimum of 1.5 years after HSCT.

Although we provide evidence that CMV related T cell expansions persist in the patient for a minimum of 1.5 years with a consistent phenotype, we cannot exclude the possibility that the expansion that is observed later on with the same TCR Vβ and a different phenotype is a similar clone that changed its phenotype over time. For instance, for the CD8 TCR Vβ4 enriched clusters, multiple clusters were detected with a different phenotype at distinct timepoints (C02-late, C12-late, C16-early, C11-early, C11-late, C62-late) in the patient and in the donor (C02-late, C59-late, Figure 7D,E, S8). Although the phenotype of the T cell expansion could change over time, after merging these clusters in one analysis, it became evident that the different intensity of CD3, CD4, and Vβ expression indicates the presence of different TCR clones with the same Vβ at day 34/d47 and a different clone at day 204/344 and in the donor (Figure S8B). The same was true for TCR Vβ14, which was highly frequent in both HD1 (Figure 2I) and HSCT1 (Figure 7E) in the late memory CD8 compartment. When analyzing the samples together, the donor and the patient clustered separately. Although there was no separation based on timepoint in the patient, the phenotypic differences suggests the existence of multiple T cell clones in the patient and a distinct clone in the healthy donor (Figure S8C).

### CMV related CD4 and CD8 T cell expansions that are donor derived still persist at 2.5 years after hematopoietic stem cell transplantation

Although in HSCT2 no CMV reactivation was observed (based on absent CMV DNA load in plasma), at day 35 post HSCT a peak in CD4 late memory (370 cells/µl) and CD8 late memory T cells (1323 cells/µl) was detected (Figure 8A,B). Neary all CD4 late memory, and CD8 T cell clusters in this patient were defined as oligoclonal expansion based on reduced TCR Vβ diversity (Figure 8C). In the CD4 and CD8 late memory T cells clusters could be defined with only one or two dominant TCR Vβs (Figure 8D,8E, S9). We selected the clusters with the highest frequency at d35 or d56 to study the kinetics of the CMV related T cell expansion after HSCT. For CD4 TCR Vβ8 and TCR Vβ13.1, cells with a similar phenotype were also present in the donor sample, while for CD4 TCR Vβ21.3 this was not the case (Figure S9). The TCR Vβ8+ cells with a TIGIT-PD-1low phenotype could be detected in HSCT2 up to d904 (Figure 8F, 8G, S10). The TCR Vβ13.1+ cells were TIGIT-PD-1+ and were also present for a minimum of 2.5 years (Figure 8H, 8I). Interestingly within the CD8 late compartment 3 TCR Vβs were enriched in clusters highly frequent at d35 that were all positive for NKG2A: Vβ9, Vβ14 and Vβ3 (Figure 8E, S10, S11A). As an example we demonstrate that the TCR Vβ9+ cells with a TIGIT+PD-1-phenotype could be detected both in the patient and donor. Two clusters (C13-HSCT2, C27-HD2) with a dominance of TCR Vβ9 and Vβ14 were excluded from the analysis because of the lower CD8 and lower side scatter, indicating dying cells (Figure S11B). The Vβ9+ cells selected from the remaining clusters with this particular TIGIT+PD-1-phenotype persisted for at least 2.5 years post HSCT (Figure 8J,K). The TCR Vβ14+ cells with the TIGIT+/-PD-1-phenotype, potentially including a TIGIT+ and TIGIT-T cell clone, were also still present at the 2.5 year timepoint (Figure 8L,M).

**Figure 8.**
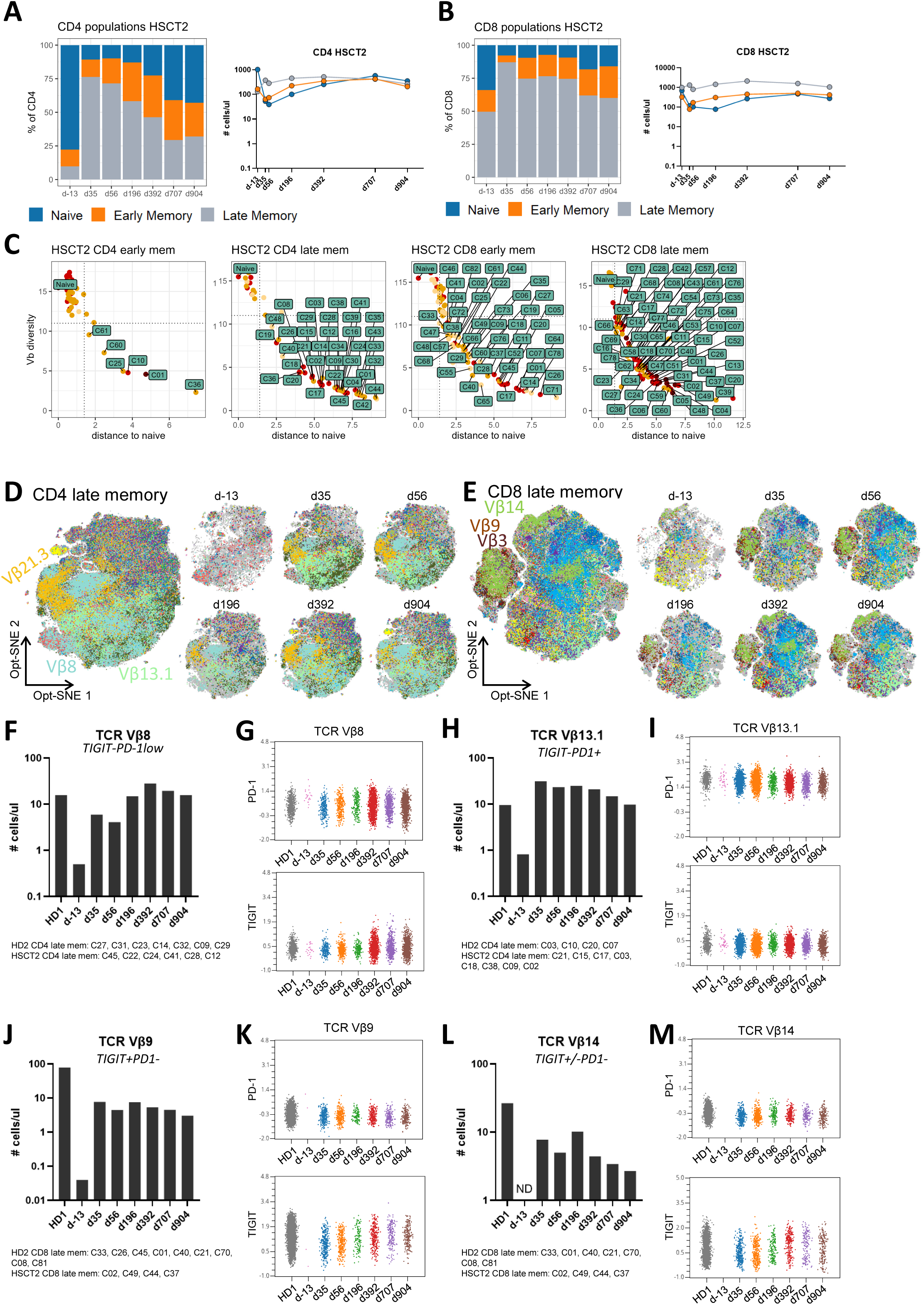
Oligoclonal T cell expansions can be traced until at least 2.5 years after HSCT in patient 2. **A.** Frequency and absolute numbers of CD4 and **B.** CD8 populations in PBMC samples before and after hematopoietic stem cell transplantation (HSCT). **C.** The samples from HSCT2 were merged and an Opt-SNE and Opt-SNE based clustering was performed on the distinct CD4 and CD8 memory populations. For every cluster the TCR Vβ diversity (inverse Simpson index) and the Euclidean distance to the naïve T cells from before the HSCT were calculated based on the frequencies of the individual Vβs per cluster. A cluster with oligoclonal expansions was defined as Vβ diversity <11, distance >1.4 and total number of Vβ+ cells >0.2% of all Vβ+ cells. The naïve cluster and the clusters that meet these criteria are labeled. **D.** Opt-SNE visualization of the CD4 late and **E.** CD8 late memory T cells. Each cell is colored by its TCR Vβ expression. **F.** From the CD4 late memory clusters with a dominance of TCR Vβ8 and highest frequency at d35/d56, the TCR Vβ8+ cells were selected and quantified. **G.** A consistent TIGIT and PD-1 (TIGIT-PD-1low) was observed over time. **H.** From the CD4 late memory clusters with a dominance of TCR Vβ13.1 and highest frequency at d35/d56, the TCR Vβ13.1+ cells were selected and quantified. **I.** A consistent TIGIT and PD-1 (TIGIT-PD-1+) was observed over time. **J.** From the CD8 late memory clusters with a dominance of TCR Vβ9 and highest frequency at d35/d56, the TCR Vβ9+ cells were selected and quantified. **K.** TIGIT and PD-1 expression per individual sample is shown for the TCR Vβ9+ cells. **L.** From the CD8 late memory clusters with a dominance of TCR Vβ14 and highest frequency at d35/d56, the TCR Vβ14+ cells were selected and quantified. The TIGIT and PD-1 phenotype is shown in **M**.

## Discussion

In this study we present a flow cytometry based method that allows deep phenotyping of T cells combined with measuring TCR Vβ expression. We developed a 9-tube 24 marker panel with a backbone panel, including antibodies to define differentiation stages and measure immune checkpoints. By combining the data from 9 tubes and perform single-cell analysis on the backbone panel of markers, we demonstrate that each T cell expansion has its own unique phenotype, that can be used to trace T cell expansions that persist for at least 2.5 years in the setting of hematopoietic stem cell transplantation.

Assessment of the TCR Vβ repertoire using flow cytometry is usually done on the bulk population of CD4 and CD8 T cells. However, it is much more valuable to study the repertoire in the context of phenotypically different T cell populations, since oligoclonal expansions can be missed. As we showed, the total CD4 and CD8 T cell TCR Vβ repertoire is reminiscent of the naïve, CM cells or CD27+CD28+ memory T cells. However, when zooming in on different memory T cell populations, oligoclonal expansions can be identified. T cells are classically subdivided into four stages of differentiation based on CCR7 and CD45RA: Naïve, CM, EM and EMRA.^26^ Previously, we observed that within one clonal T cell expansion CD45RA is heterogeneously expressed, making CD45RA not useful to capture oligoclonal expansions.^22^ Therefore we employed a different strategy in which we categorized memory T cells into early and late based on the CD27 and CD28. Indeed, most oligoclonal expansions were identified either in the early memory T cells (CD27+CD28+ T cells) or in the late memory T cells (CD27-and/orCD28-).

In the memory CD4 and CD8 T cell compartment, the KLRG1+ cells had reduced TCR Vβ diversity. KLRG1, an inhibitory receptor, has been previously reported to be expressed by highly cytotoxic effector CD8 T cells in mice.^27–29^ Since these KLRG1 effector CD8 T cells are short-lived, this raises the question how the pool of memory cells is maintained. Longitudinal tracking of KLRG1+ effector CD8+ T cells in vivo showed that they displayed developmental plasticity, since they were able to downregulate KLRG1 and enter the pool of long-lived KLRG1– CD127+ memory cells.^30^ Graded CX3CR1 expression, together with CD62L, has been reported in both human and mice to mark the T cell differentiation continuum.^31^ In line with this, we observed the lowest TCR Vβ diversity in the late memory KLRG1+CX3CR1+ population.

By clustering analysis we identified clusters in which only one or two TCR Vβ were dominant, mostly in the CD4 and CD8 late compartment. Of note, these cells clustered based on their phenotype and not the TCR Vβ itself, since the TCR Vβs were not included in the clustering. Therefore, it’s the repertoire of surface receptors that drove the clustering. On the CD4 and CD8 T cells, the TIGIT and PD-1 expression was most important in driving the clustering, while CD57, CD56 and CD45RA were heterogeneously expressed. NKG2A was only observed on a fraction of the CD8 memory T cells. Of note, CD3+CD56+ cells are usually excluded from the conventional T cells and labeled as NKT cell. Here we provide evidence, that within a T cell expansion CD56 can be bimodally expressed, reinforcing the need to include the CD3+CD56+ cell in the T cell gate. The same holds true for CD4+CD8dim cells, since on a fraction of cells within certain CD4 T cell expansions CD8 expression was detected.

To monitor T cells expansions we longitudinally studied the circulating T cell compartment in two hematopoietic stem cell transplant (HSCT) recipients for which the patient and donor were CMV seropositive. Although there was no CMV reactivation in the second patient, still a CD8 T cell expansion was observed. In agreement with the literature, if the donor is CMV seropositive the CD8 T cells expand, even in the absence of reactivation.^32^ Using our single-cell analysis method we were able to trace CMV associated CD4 and CD8 T cell expansions that mirrored populations present in the donor. It has been reported that up to 1 year after HSCT the CMV-specific response mainly relies on donor mature T cells in the graft.^33^ Although we cannot exclude the possibility that the same clone evolves over time with a different phenotype, we do provide evidence that the same oligoclonal expansion can exist for over a period of 2.5 years with a consistent TIGIT and PD-1 phenotype.

Overall, we demonstrate that a flow cytometry assay combined with single-cell analysis is a useful method to monitor oligoclonal T cell expansions. Our method is widely applicable and can be used as diagnostic tool study responses in infection, vaccination, autoimmunity, transplantation and tumor immunology. More efficient use of TCR Vβ antibodies by increasing the number of conjugated fluorochromes will reduce the number of tubes required and thereby limit the material that is needed to run the assay. Understanding memory formation of T cells and their persistence is of considerable importance in future strategies which aim to enhance T cell immunity, especially in the context of immune checkpoint inhibitors.

