## Supplementary material for "Immunophenotyping of T cells Combined with Vβ antibodies identifies long lasting CMV related T cell Expansions with a consistent TIGIT and PD-1 phenotype": Sup figure

### Figure S1

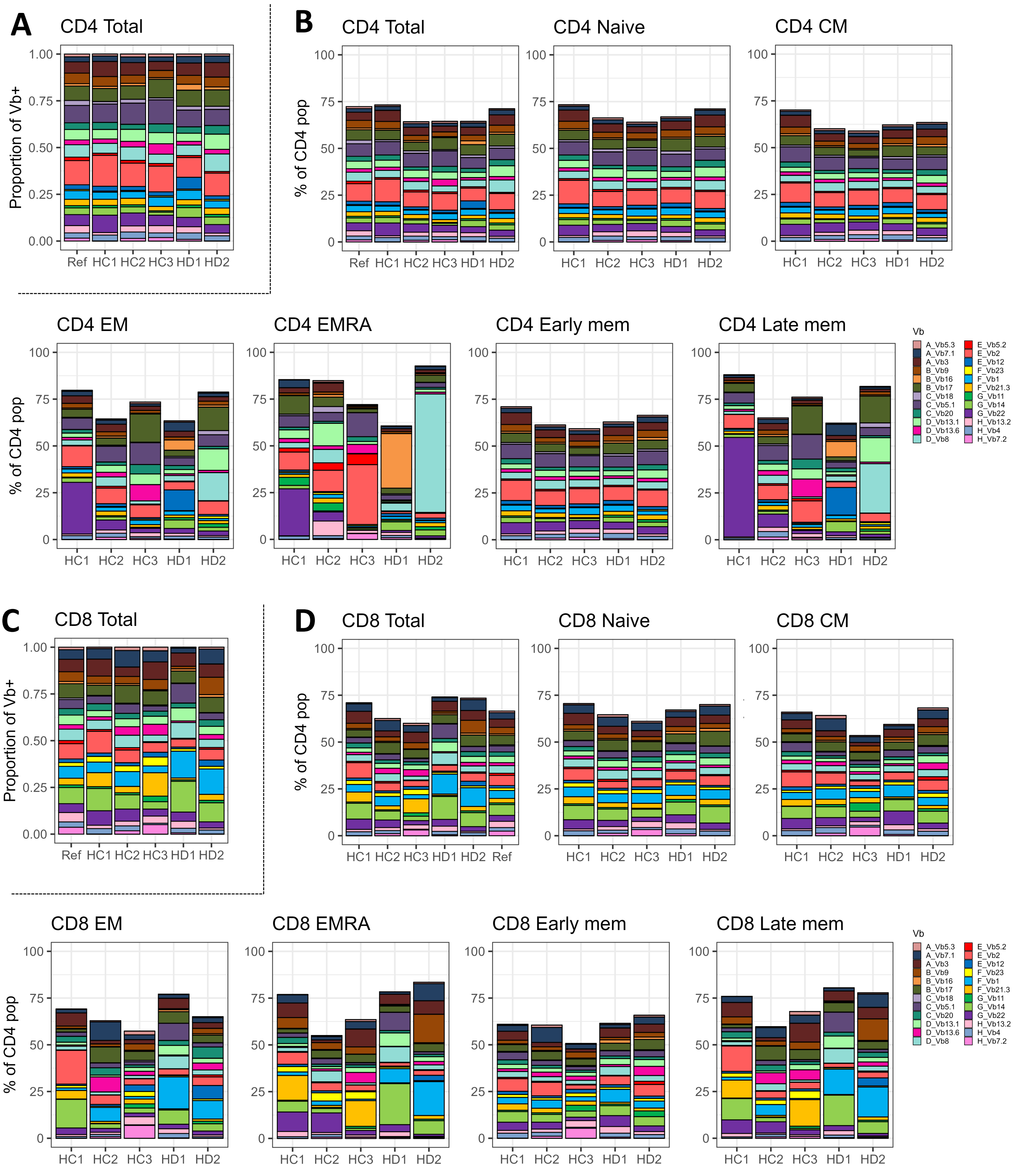


**Figure S1. TCR Vβ distribution in healthy donors per T cell population**

**A.** The TCR Vβ distribution per individual donor, for the total CD4 T cells. As a reference (ref), the mean percentage of total CD4 T cells for every single TCR Vβ provided by the Vβ antibody kit is shown. **B.** The percentage of CD4 T cells that was stained by one of the Vβ antibodies. The 24 Vβ antibodies cover approximately 70% of the TCR Vβ repertoire. Dependent on clonal expansions with a certain TCR Vβ that is covered or not covered by the kit, this percentage could be a bit higher or lower in the memory populations. **C.** The TCR Vβ distribution per individual donor, for the total CD8 T cells. As a reference, the mean percentage of total CD8 T cells for every single TCR Vβ provided by the Vβ antibody kit is shown. **D.** The percentage of CD8 T cell populations that was stained by one of the Vβ antibodies. HC = healthy control, HD = healthy donor.

### Figure S2 CD4 early memory

## HC1

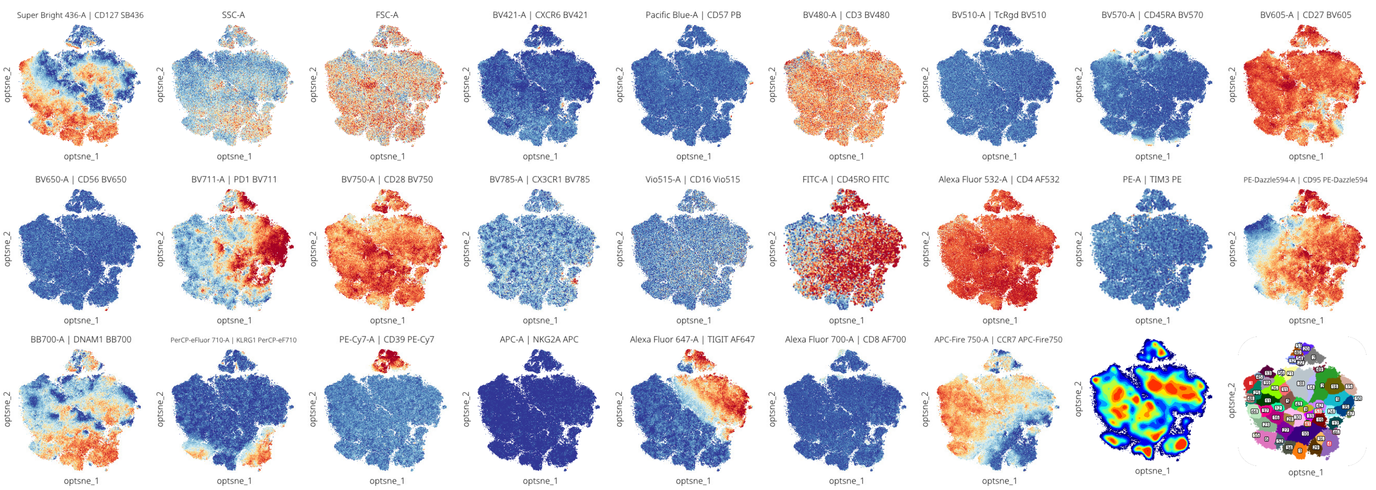


## HC2

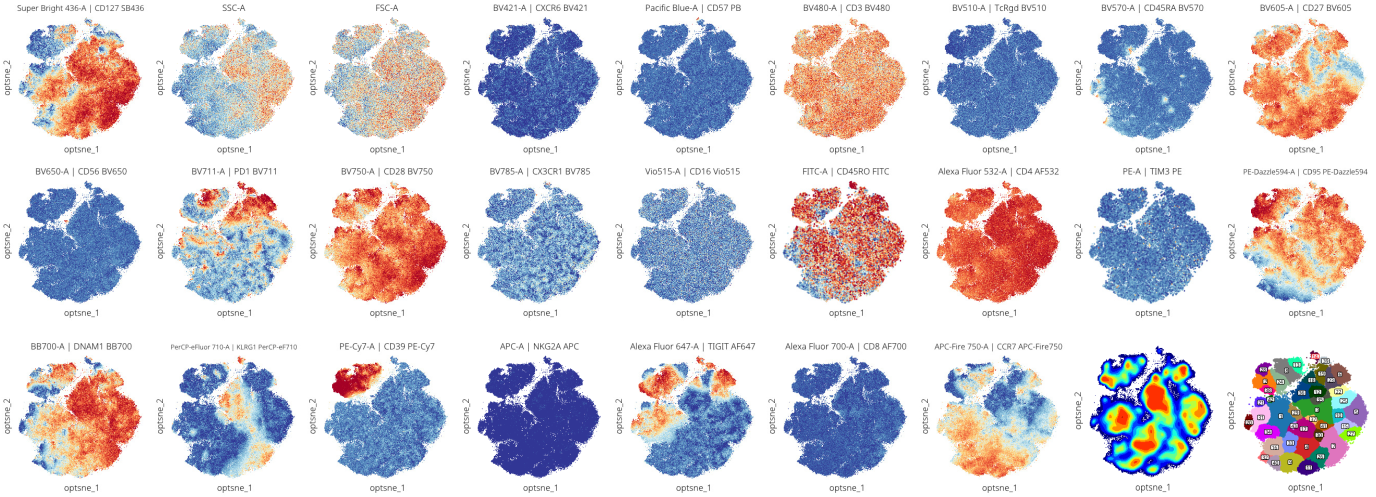


## HC3

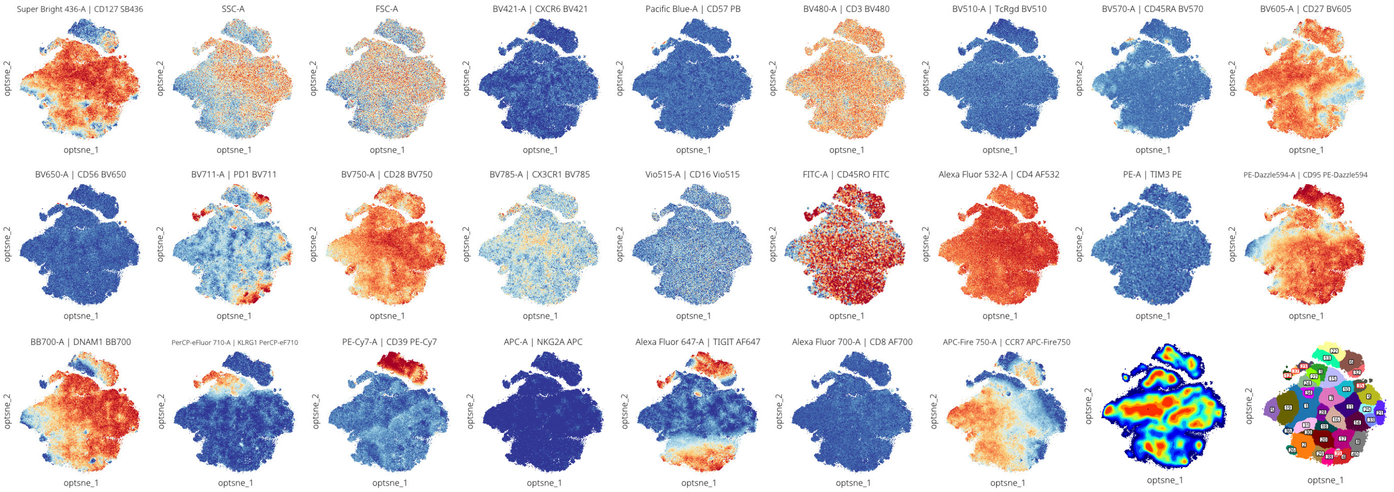

### Figure S2 CD4 early memory

## HD1

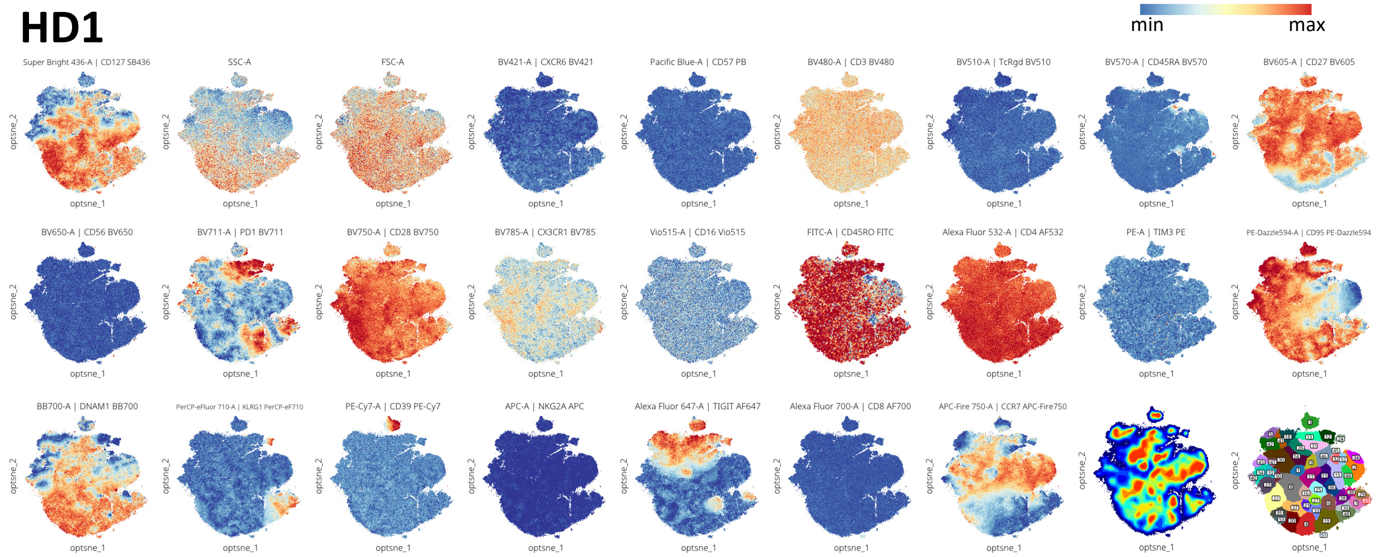


## HD2

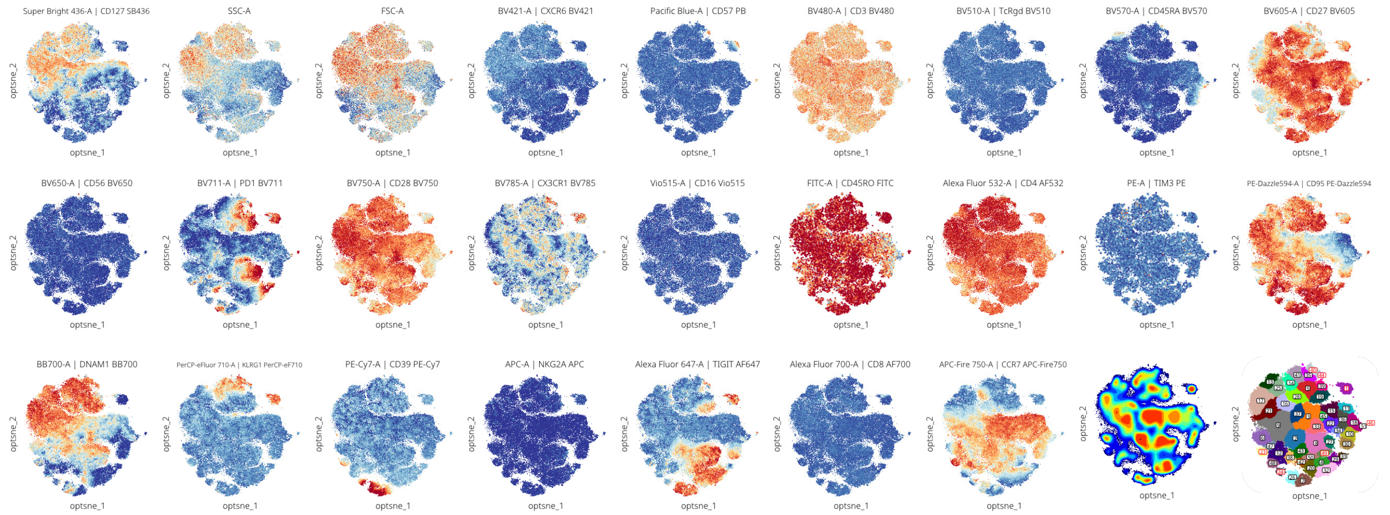


**Figure S2. Expression of markers on early memory CD4 T cells per individual donor**
Opt-SNE embedding was based on all fluorescent markers except CD3, TCRgd, CD16, CD4, CD45RO, TIM-3 and Vβs. The nine files per healthy control (HC) or donor (HD) were combined for the Opt-SNE. For the CD45RO and TIM-3 expression only cells from tube 9 are shown. The density-based clustering was done by ClusterX based on the Opt-SNE coordinates. The clusters that were defined as oligoclonal expansion in Figure 3B, are labeled in red.

### Figure S3 CD4 late memory

## HC1

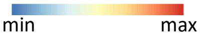


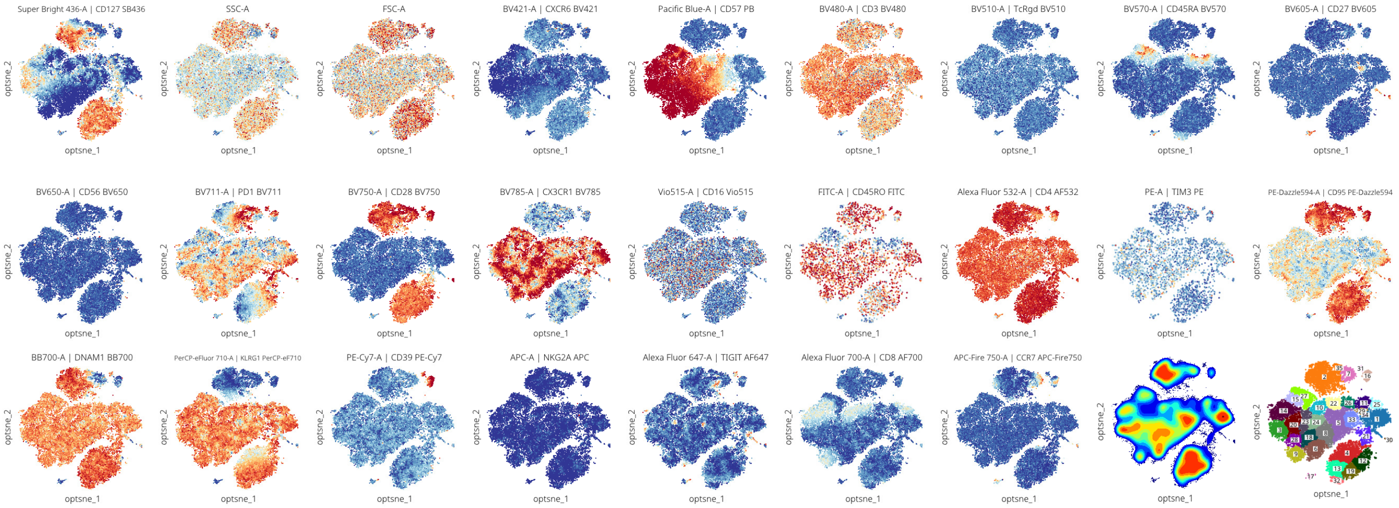


## HC2

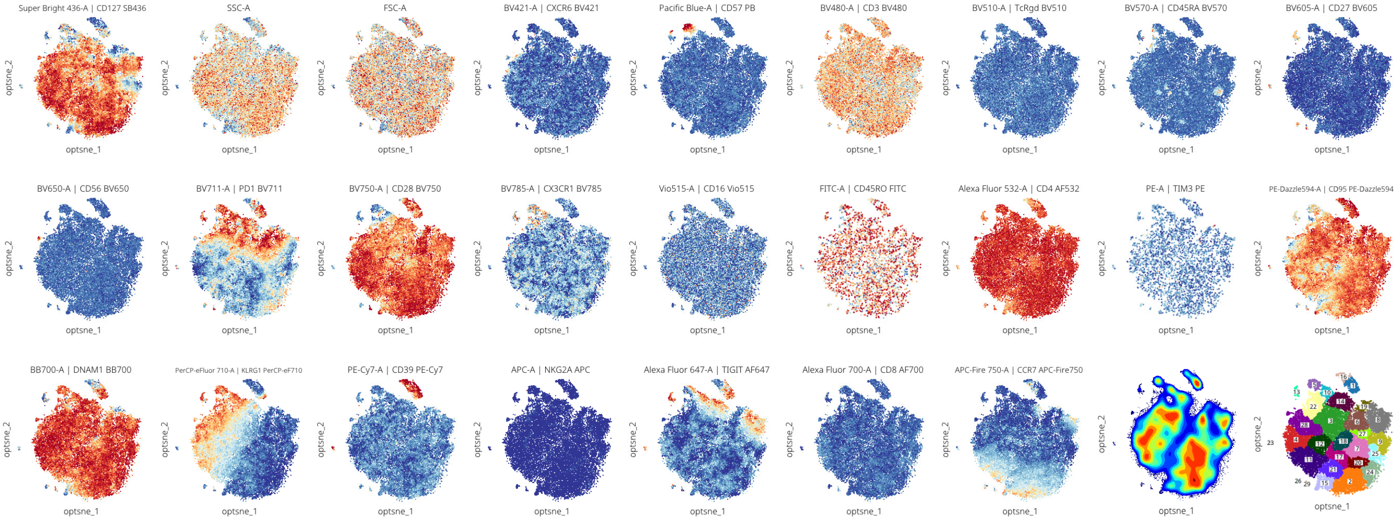


## HC3

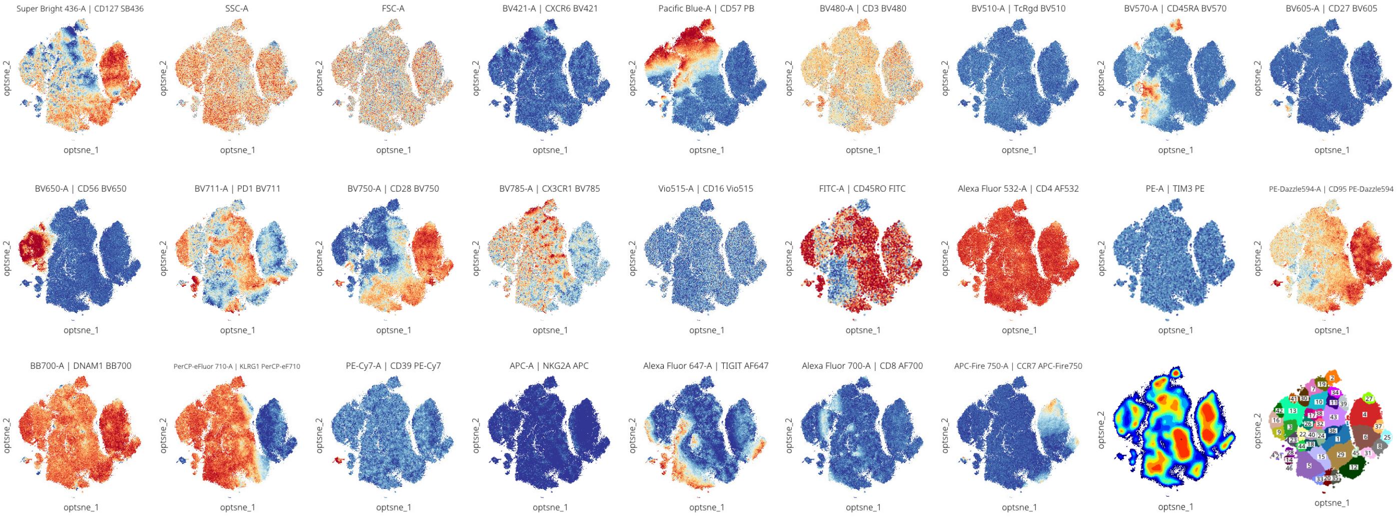

### Figure S3 CD4 late memory

## HD1

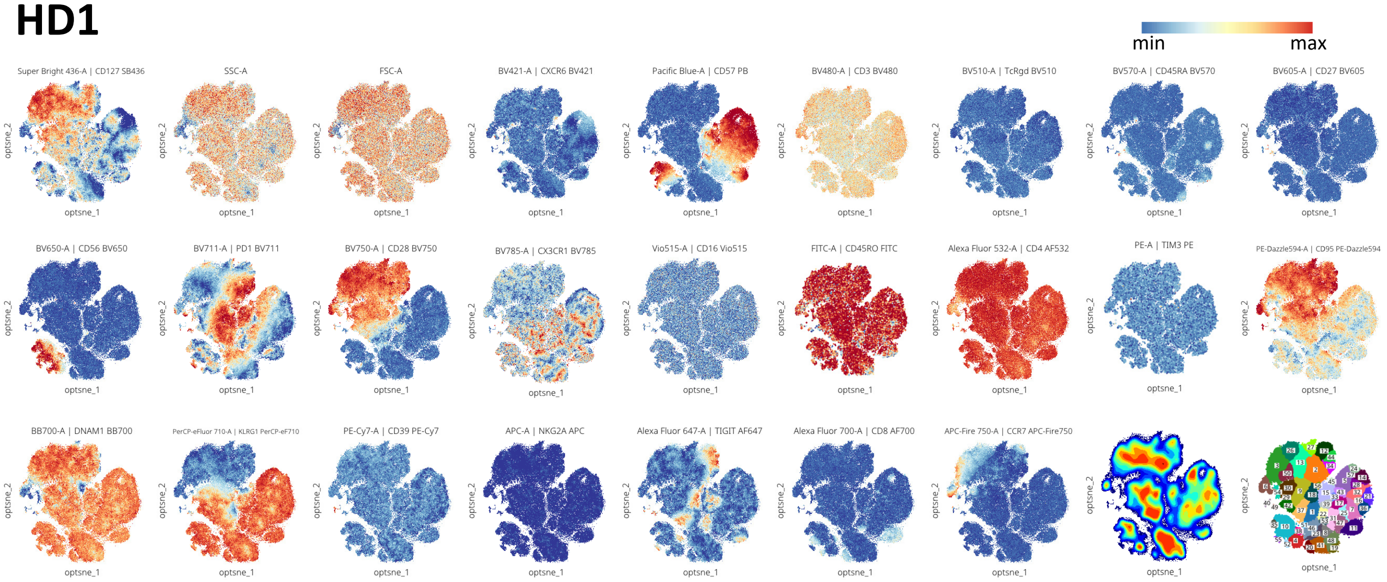


## HD2

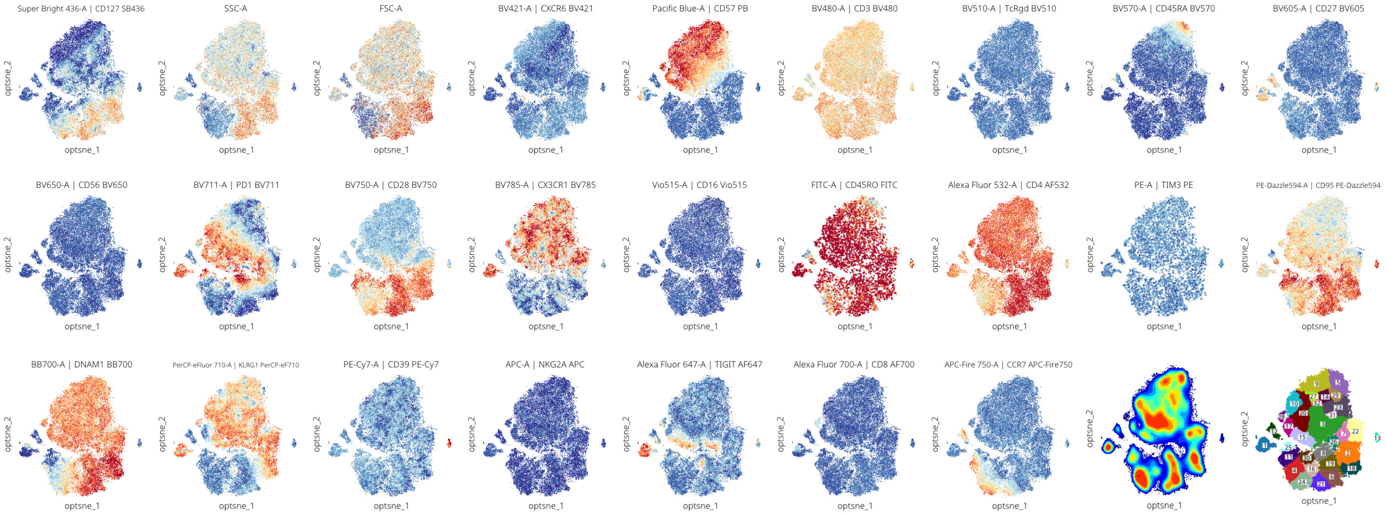


**Figure S3. Expression of markers on late memory CD4 T cells per individual donor**
Opt-SNE embedding was based on all fluorescent markers except CD3, TCRgd, CD16, CD4, CD45RO, TIM-3 and Vβs. The nine files per healthy control (HC) or donor (HD) were combined for the Opt-SNE. For the CD45RO and TIM-3 expression only cells from tube 9 are shown. The density-based clustering was done by ClusterX based on the Opt-SNE coordinates.

### Figure S4 CD8 early memory

## HC1

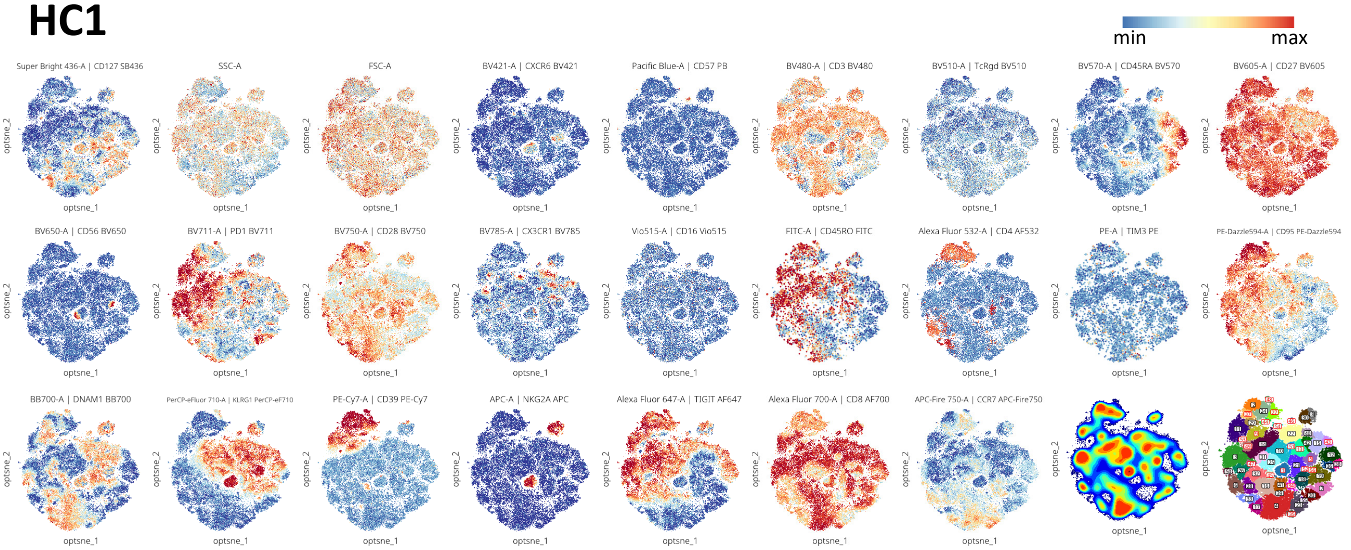


## HC2

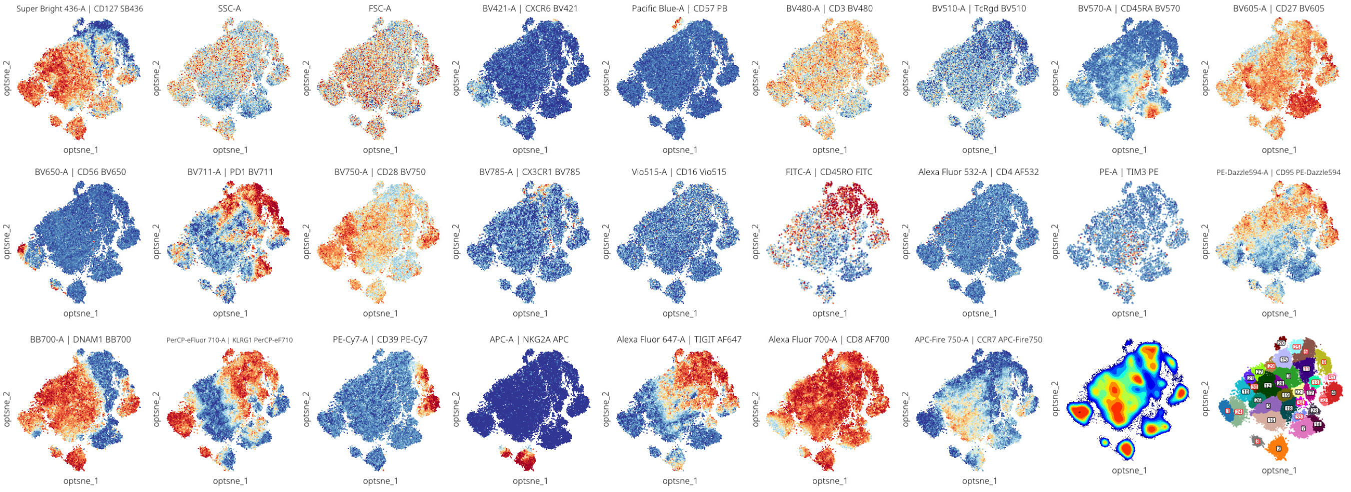


## HC3

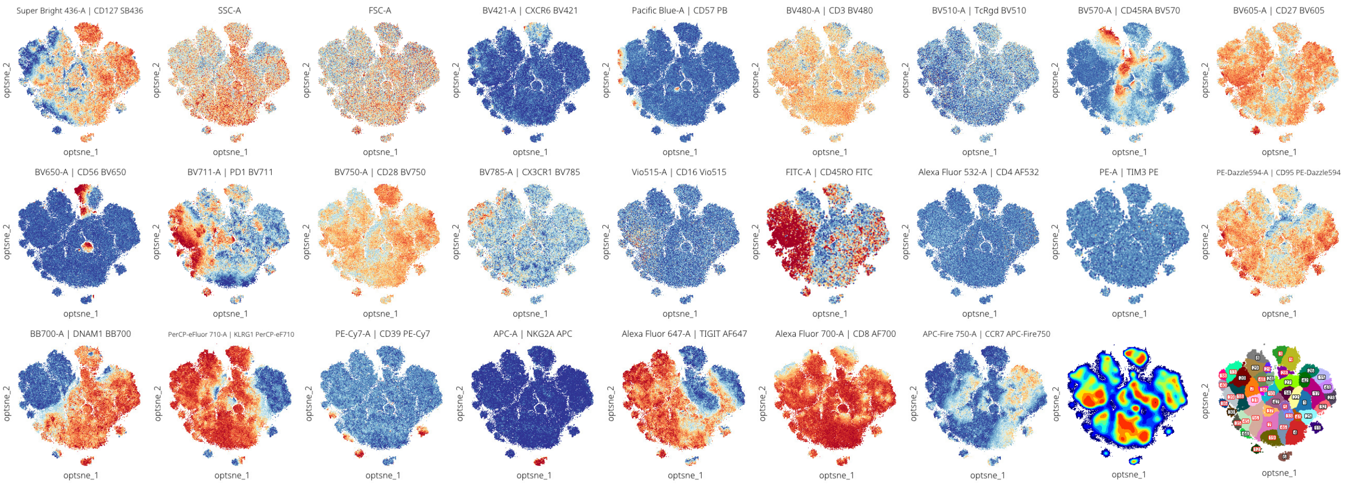

### Figure S4 CD8 early memory

## HD1

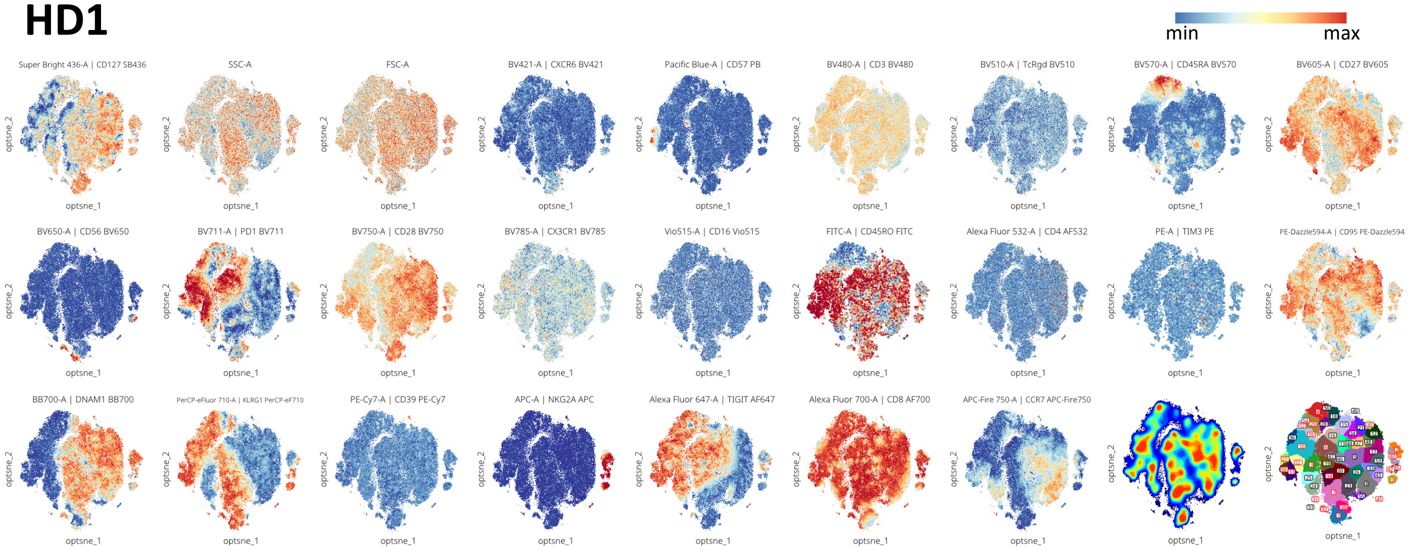


## HD2

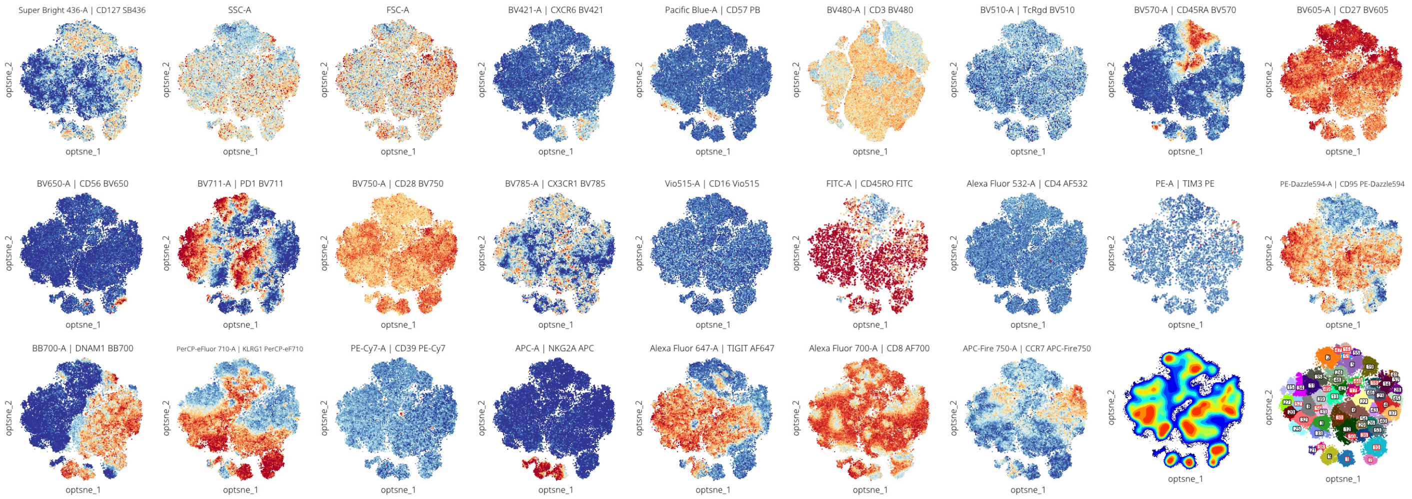


**Figure S4. Expression of markers on early memory CD8 T cells per individual donor**
Opt-SNE embedding was based on all fluorescent markers except CD3, TCRgd, CD16, CD4, CD45RO, TIM-3 and Vβs. The nine files per healthy control (HC) or donor (HD) were combined for the Opt-SNE. For the CD45RO and TIM-3 expression only cells from tube 9 are shown. The density-based clustering was done by ClusterX based on the Opt-SNE coordinates. The clusters that were defined as oligoclonal expansion in Figure 5B, are labeled in red.

### Figure S5 CD8 late memory

## HC1

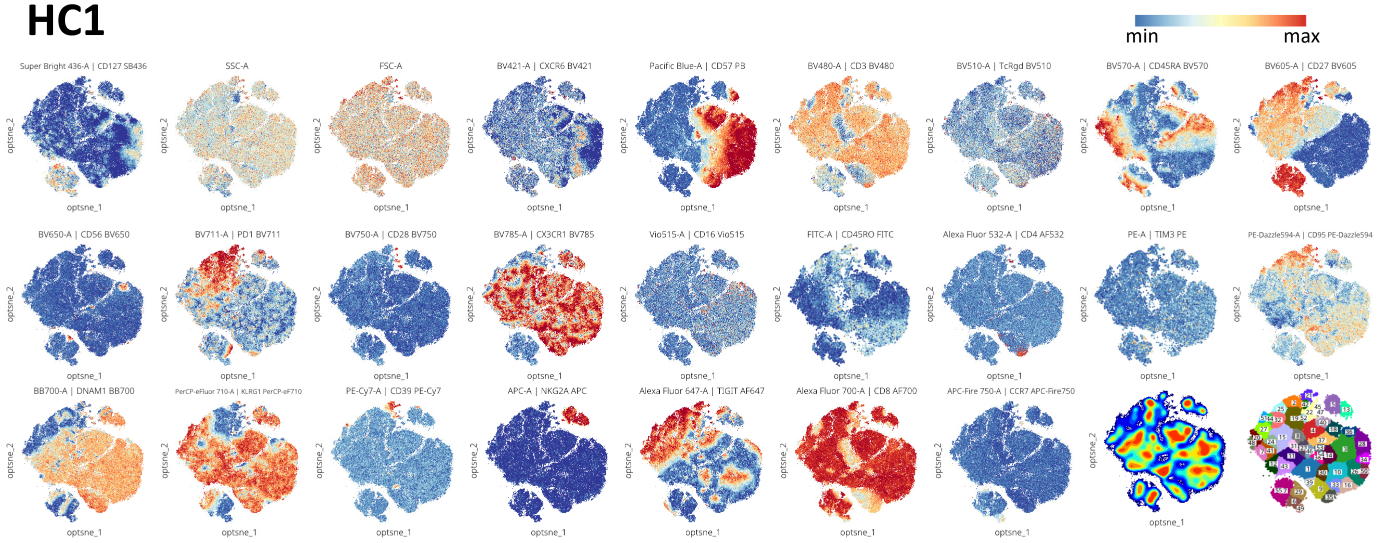


## HC2

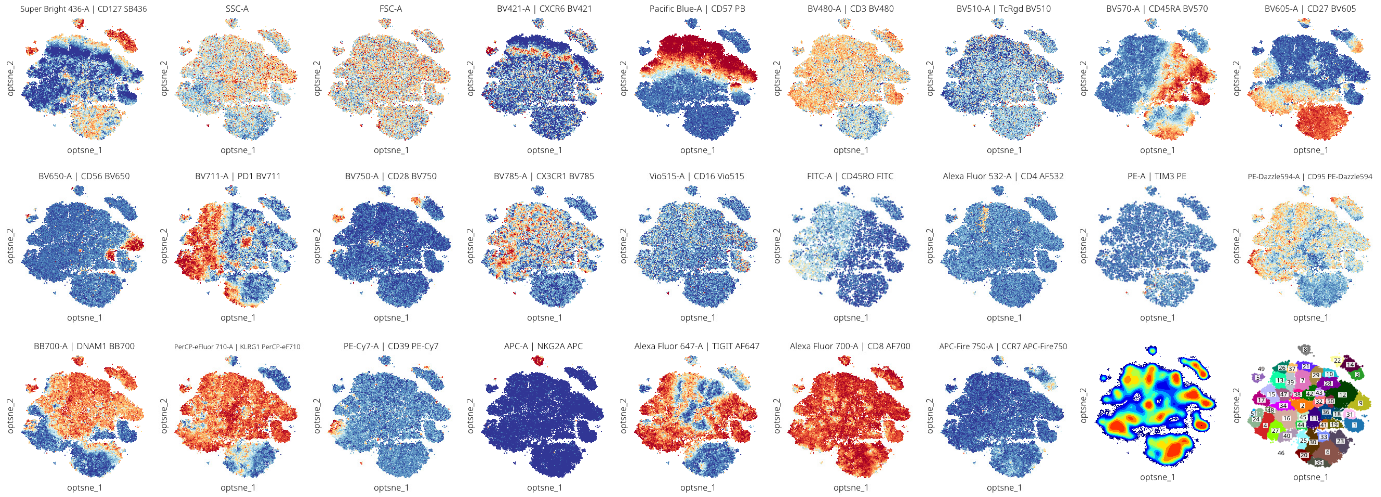


## HC3

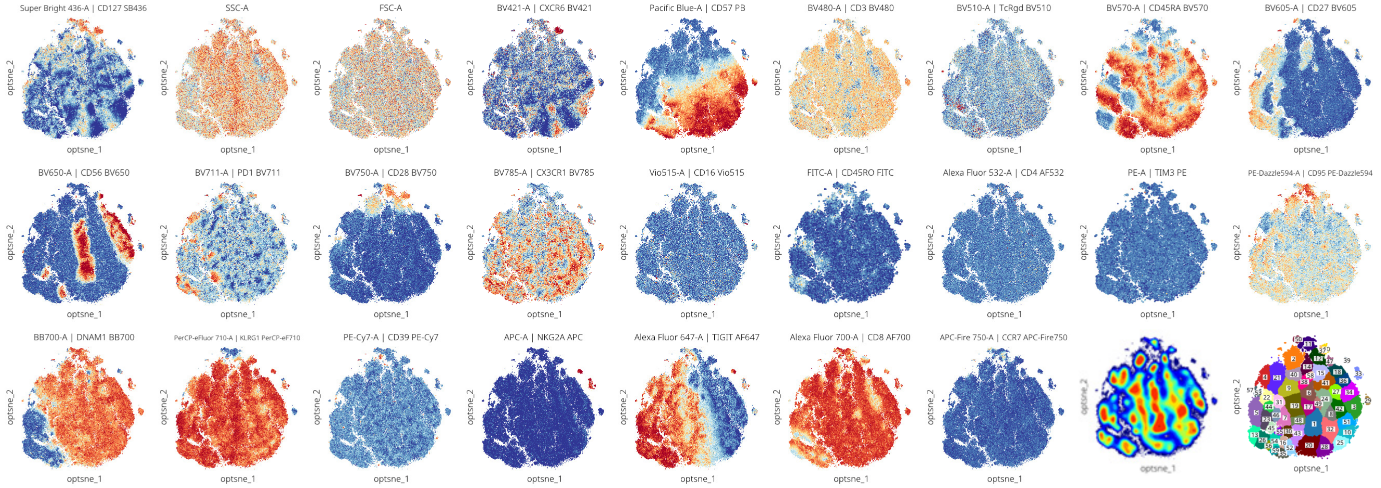

### Figure S5 CD8 late memory

## HD1

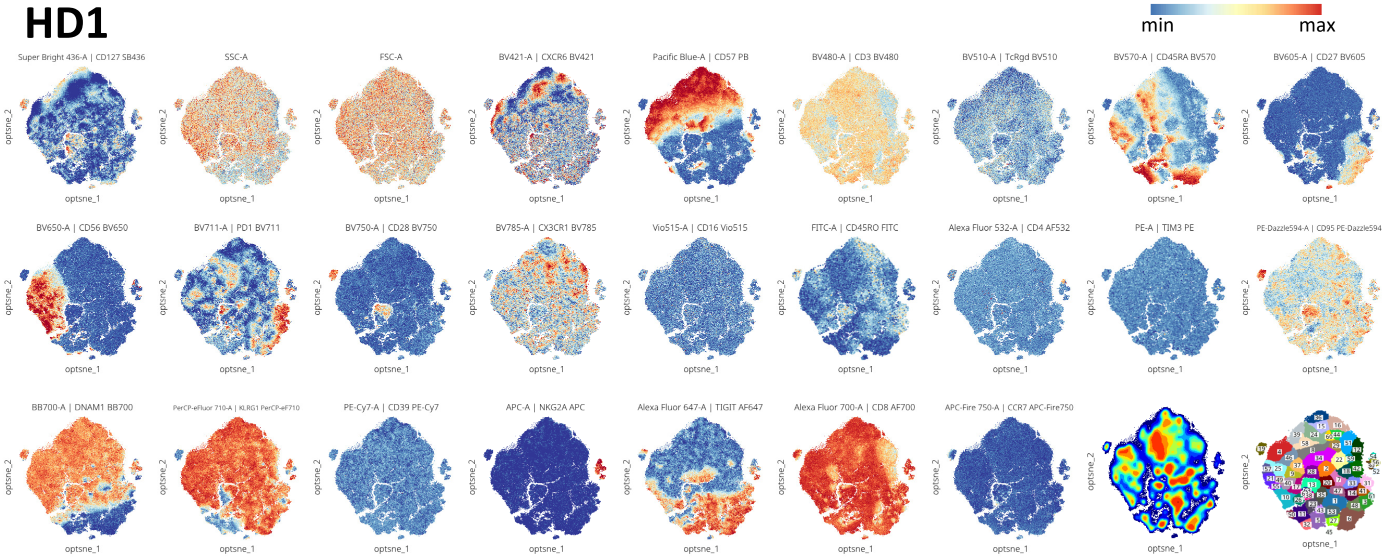


## HD2

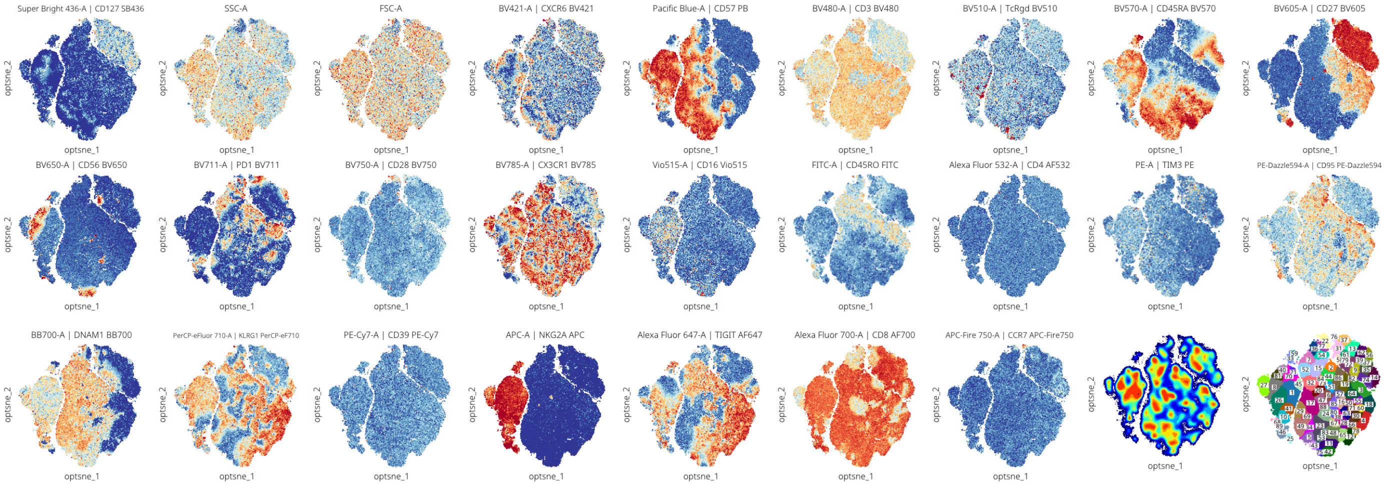


**Figure S5. Expression of markers on late memory CD8 T cells per individual donor**
Opt-SNE embedding was based on all fluorescent markers except CD3, TCRgd, CD16, CD4, CD45RO, TIM-3 and Vβs. The nine files per healthy control (HC) or donor (HD) were combined for the Opt-SNE. For the CD45RO and TIM-3 expression only cells from tube 9 are shown. The density-based clustering was done by ClusterX based on the Opt-SNE coordinates.

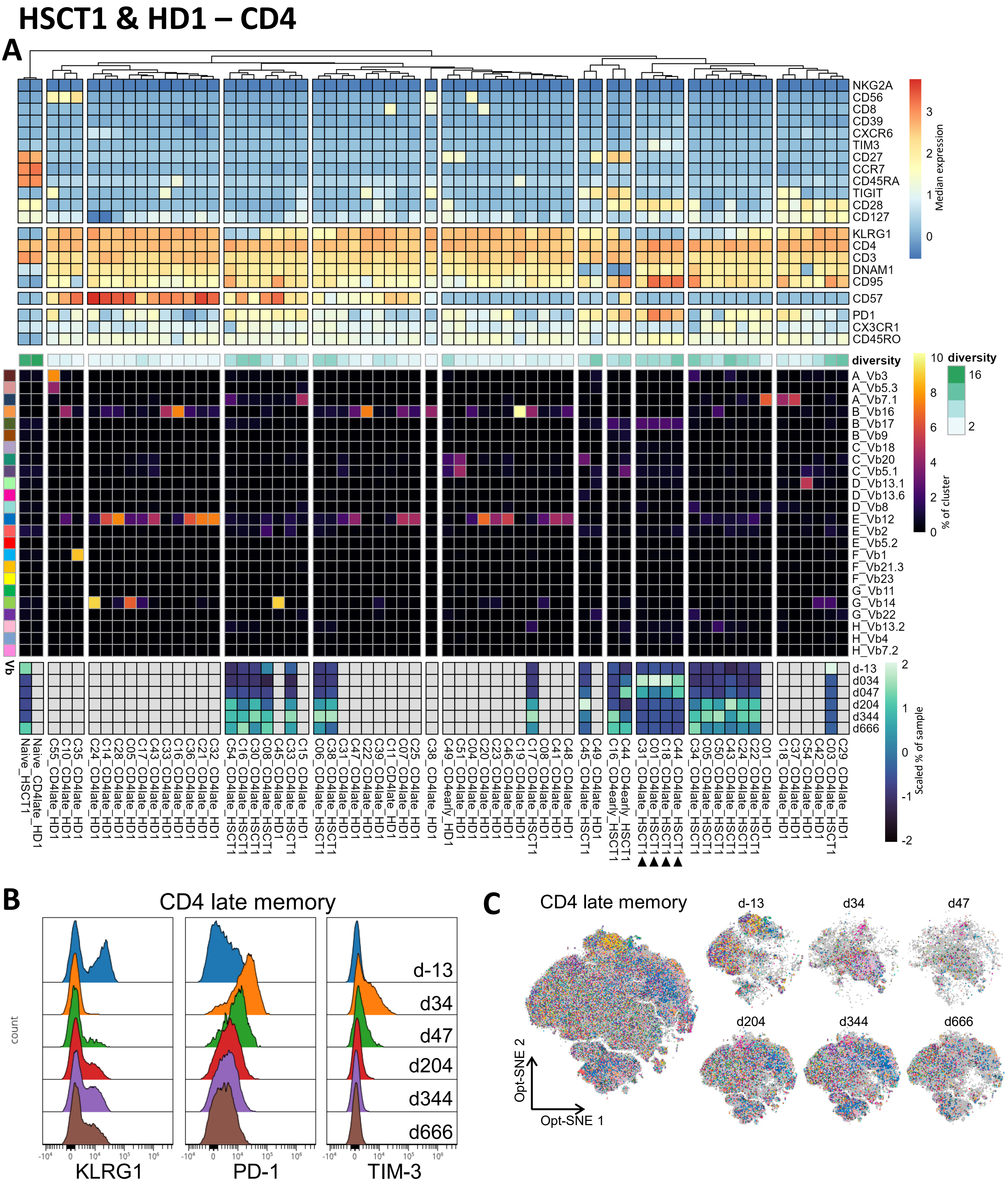


**Figure S6. Phenotype and frequency of Vβ dominant CD4 clusters of both patient HSCT1 and its donor HD1**

**A.** The CD4 early and late memory clusters that were defined as oligoclonal expansions in Figure 3, 4 and 7 are included in the heatmap. The top heatmap represents the phenotype (median intensity), the second the TCR Vβ diversity (inverse Simpson index), the third the Vβ frequency and the bottom the scaled cluster frequency for the individual HSCT1 samples. As a reference the naïve CD4 T cells are included. The arrowheads point to the clusters as discussed in the text. **B.** The expression of KLRG1, PD-1 and TIM-3 is shown on the CD4 late memory T cells from HSCT1. **C.** Opt-SNE visualization of the CD4 late memory T cells. Each cell is colored by its TCR Vβ expression.

### Figure S7 HSCT1 patient

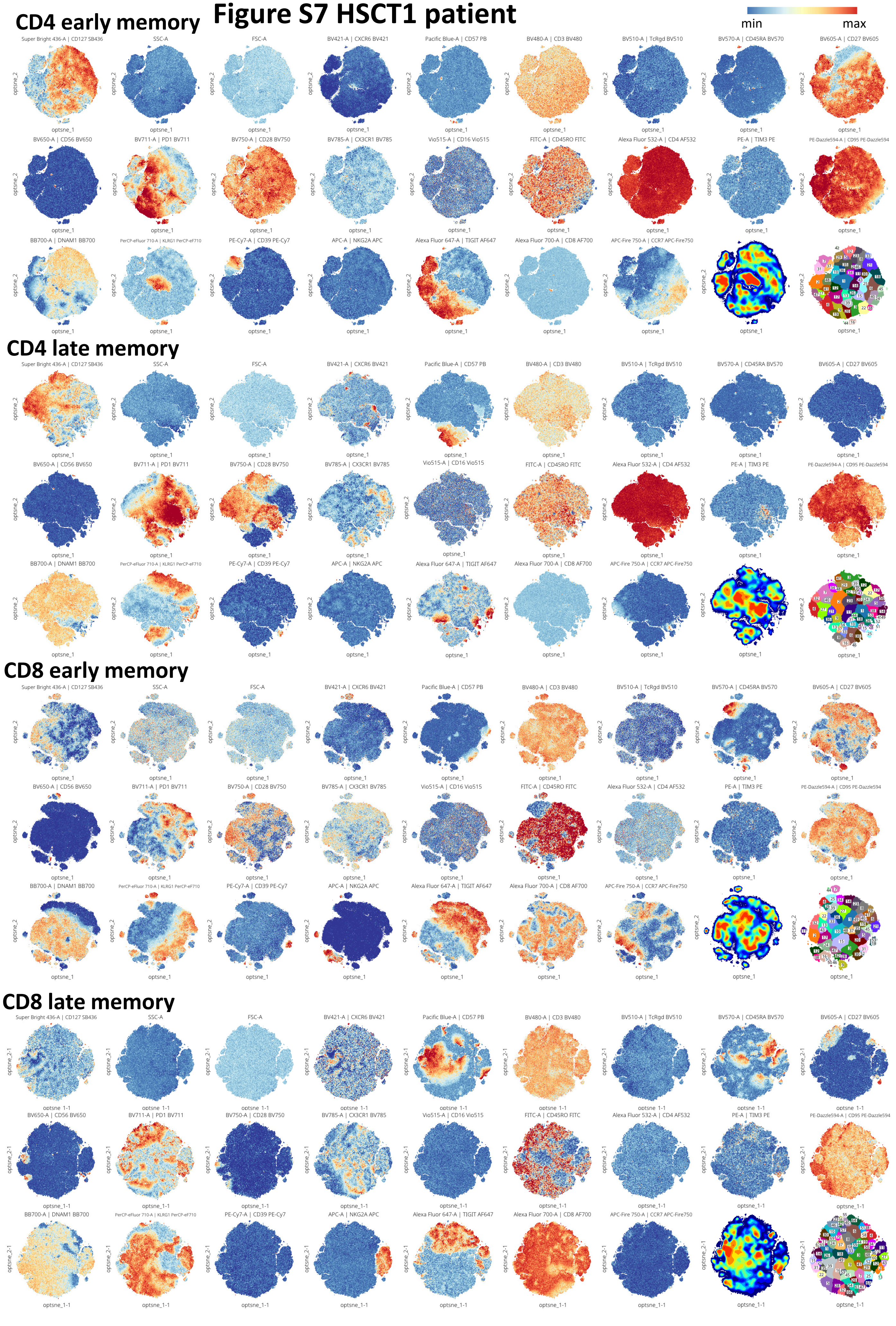

**Figure S7. Expression of markers on T cell populations from HSCT1**

Opt-SNE embedding was based on all fluorescent markers except CD3, TCRgd, CD16, CD4, CD45RO, TIM-3 and Vβs. The 6 files from different timepoints from HSCT1 were combined for the Opt-SNE. For the CD45RO and TIM-3 expression only cells from tube 9 are shown. The density-based clustering was done by ClusterX based on the Opt-SNE coordinates.

### HSCT1 & HD1 – CD8

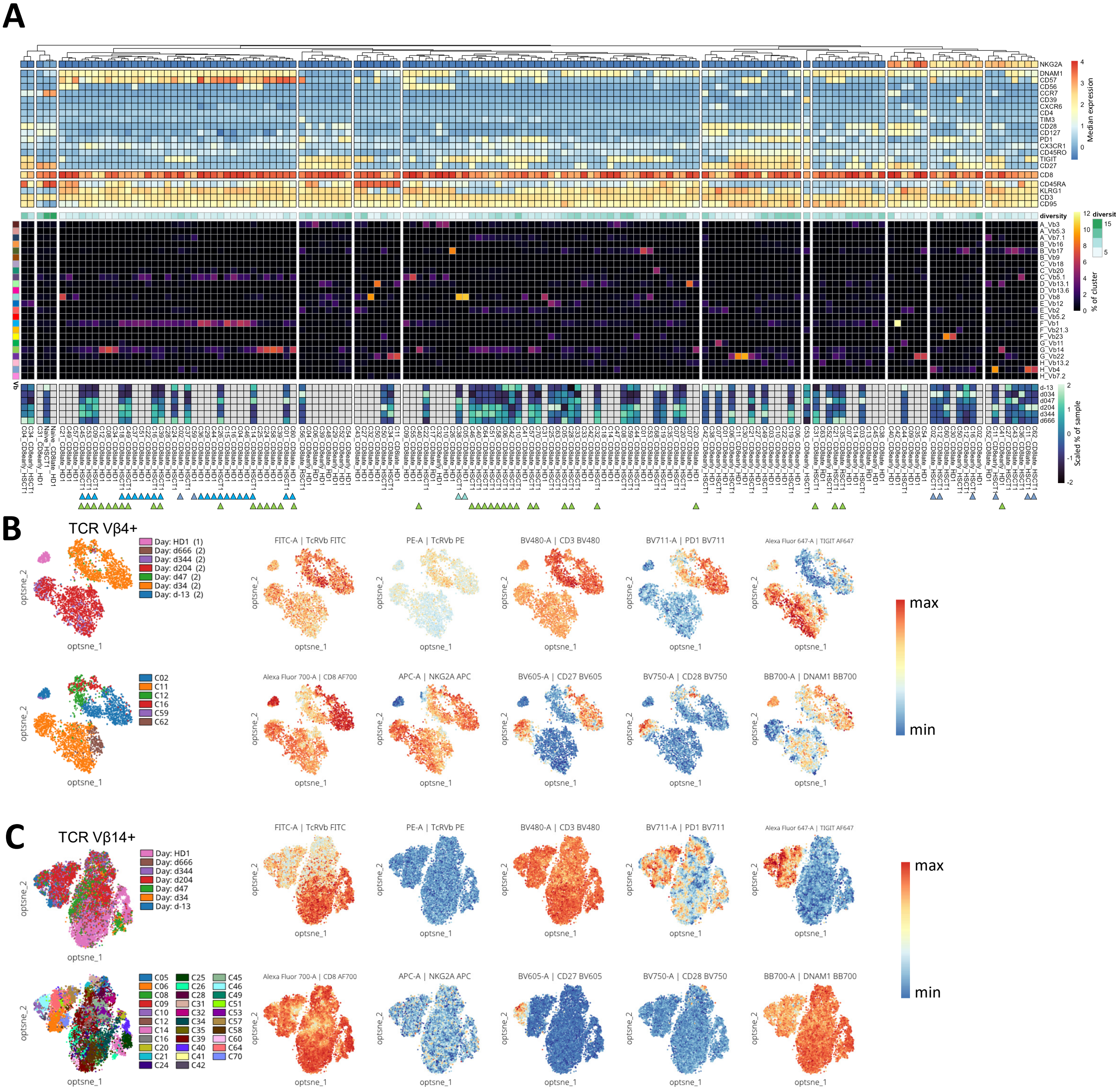


**Figure S8. Phenotype and frequency of Vβ dominant CD8 clusters of both patient HSCT1 and its donor HD1**

**A.** The CD8 early and late memory clusters that were defined as oligoclonal expansions in Figure 5, 6 and 7 are included in the heatmap. The top heatmap represents the phenotype (median intensity), the second the TCR Vβ diversity (inverse Simpson index), the third the Vβ frequency and the bottom the scaled cluster frequency for the individual HSCT1 samples. As a reference the naïve CD4 T cells are included. The arrowheads point to the clusters that were selected for further quantification and characterization in Figure 7F-I, panel B or C. The color represents the TCR Vβ. **B** TCR Vβ4 and enriched clusters from HSCT (C11-early, C16-early, C11-late, C62-late, C02-late, C12-late) and HD1 (C02-late, C59-late) were merged and an Opt-SNE was performed on the TCR Vβ4+ cells. The distinct intensity of TCR Vβ expression points towards the presence of different T cell clones in the donor and at day 34/47 and day 204/344. **C.** TCR Vβ14 enriched clusters from HSCT1 and HD1 were merged and an Opt-SNE was performed on the TCR Vβ14+ cells. The distinct intensity of the TCR Vβ suggests that the TCR clone differs between the patient and the donor. Opt-SNE in B and C was based on the backbone panel of markers, excluding CD3, TCRgd, CD16, and CD4.

### HSCT2 & HD2 – CD4

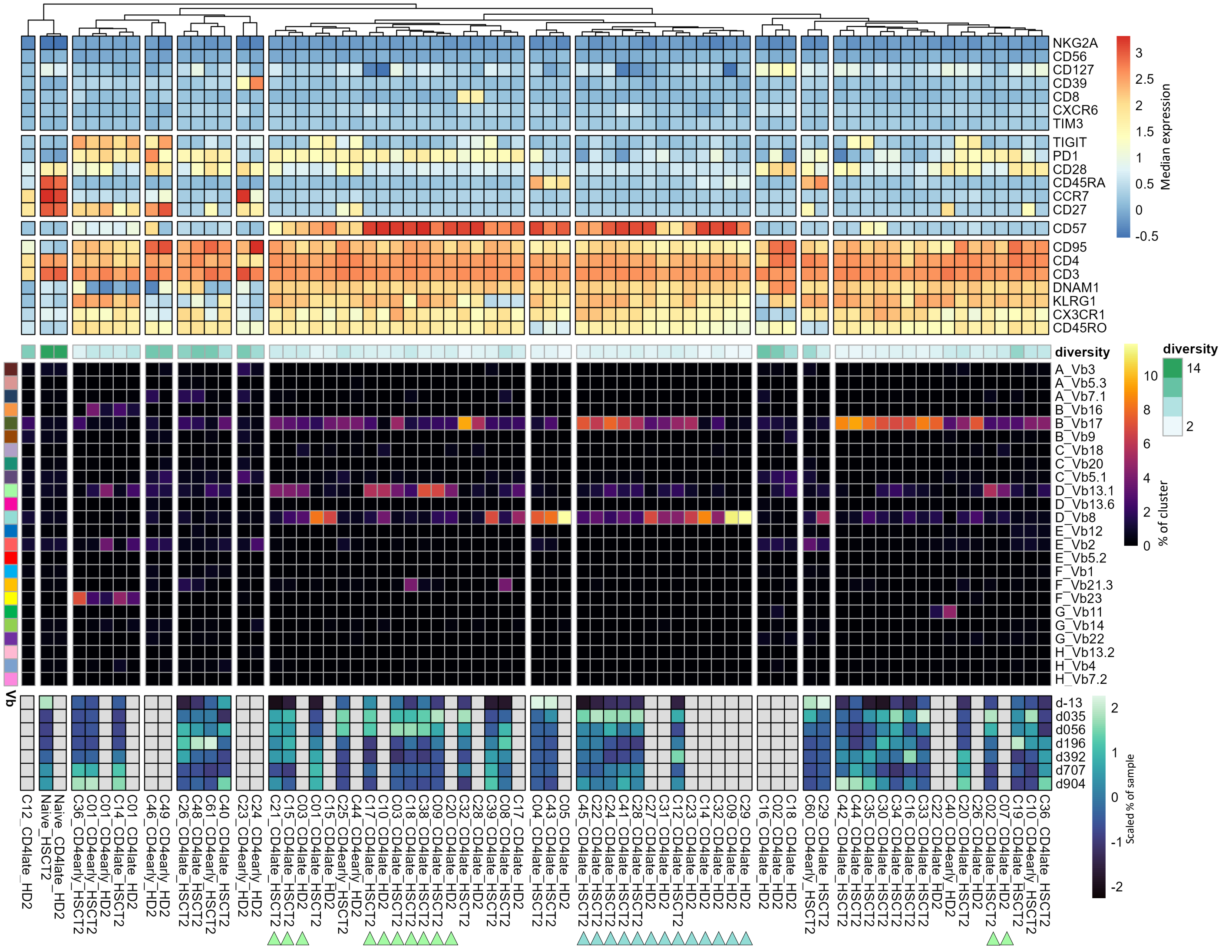

**Figure S9. Phenotype and frequency of Vβ dominant CD4 clusters of both patient HSCT2 and its donor HD2**

The CD4 early and late memory clusters that were defined as oligoclonal expansions in Figure 3 (HD2), 4 (HD2) and 8 (HSCT2) are included in the heatmap. The top heatmap represents the phenotype (median intensity), the second the TCR Vβ diversity (inverse Simpson index), the third the Vβ frequency and the bottom the scaled cluster frequency for the individual HSCT2 samples. As a reference the naïve CD4 T cells are included. The arrowheads point to the clusters that are further quantified and characterized in Figure 8F-I. The color represents the TCR Vβ

### Figure S10 HSCT2

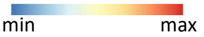


#### CD4 early memory

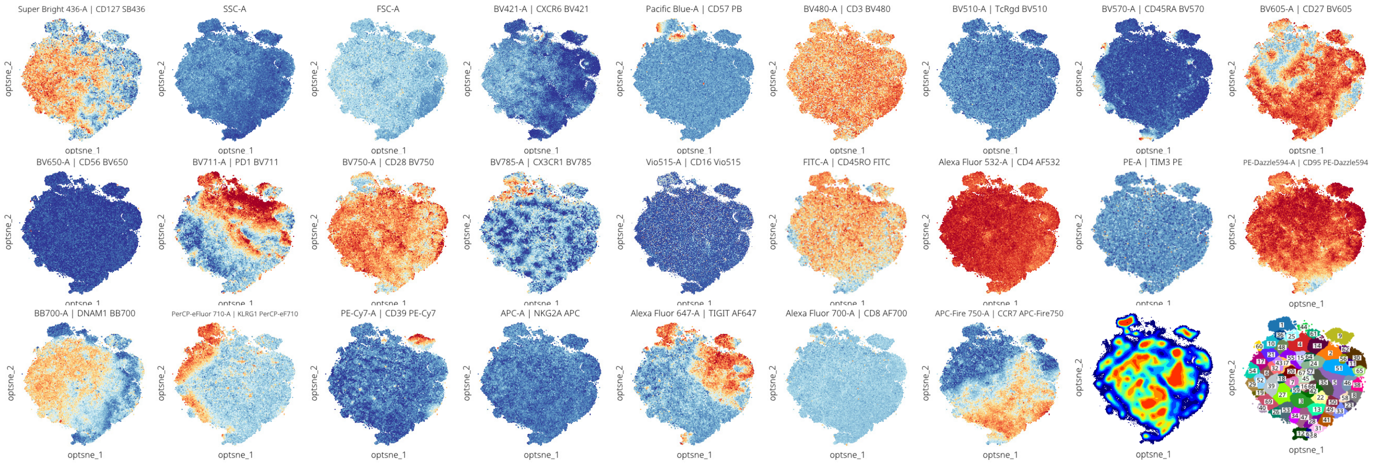


#### CD4 late memory

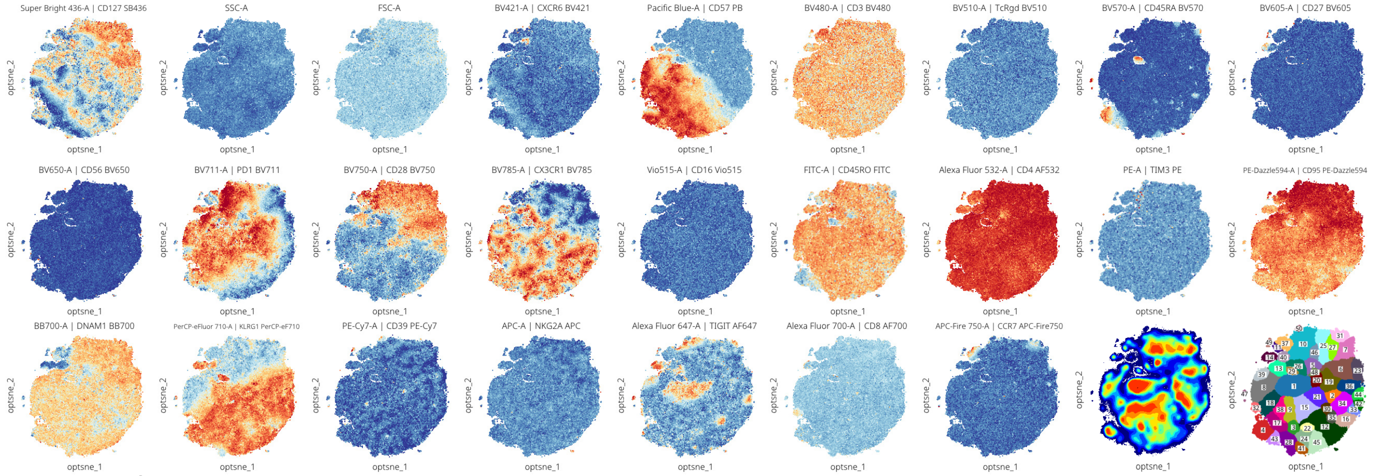


#### CD8 early memory

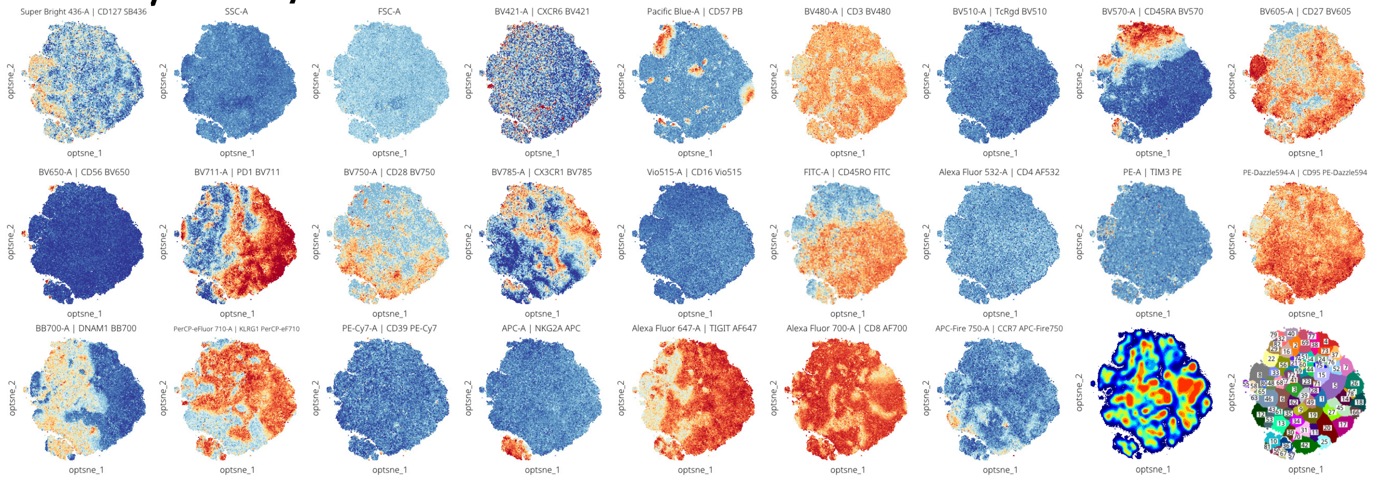


#### CD8 late memory

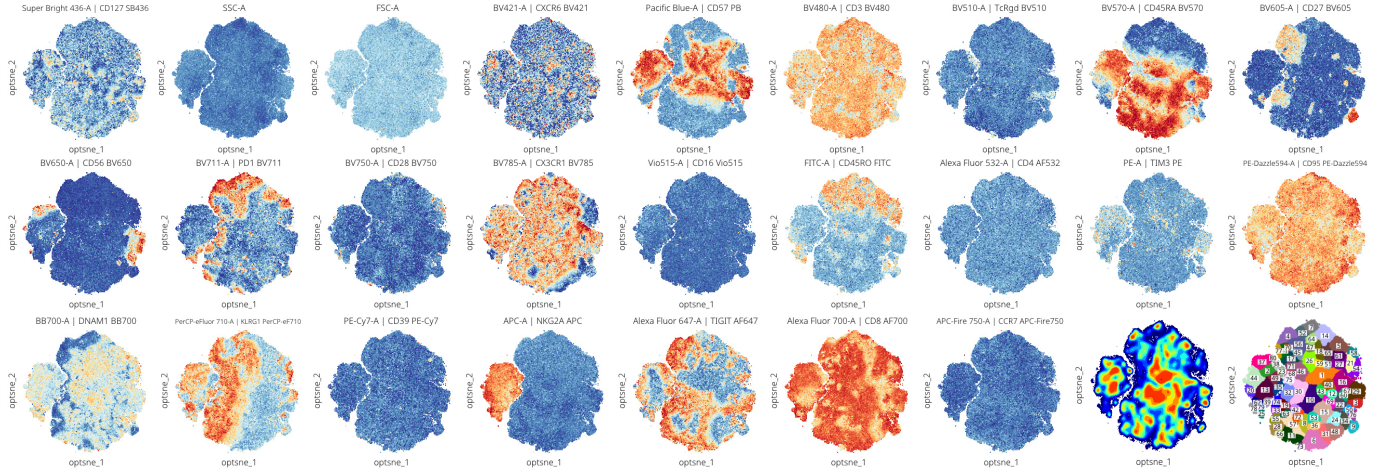

**Figure S10. Expression of markers on T cell populations from HSCT1**
Opt-SNE embedding was based on all fluorescent markers except CD3, TCRgd, CD16, CD4, CD45RO, TIM-3 and Vβs. The 6 files from different timepoints from HSCT2 were combined per population for the Opt-SNE. For the CD45RO and TIM-3 expression only cells from tube 9 are shown. The density-based clustering was done by ClusterX based on the Opt-SNE coordinates.

### HSCT2 & HD2 – CD8

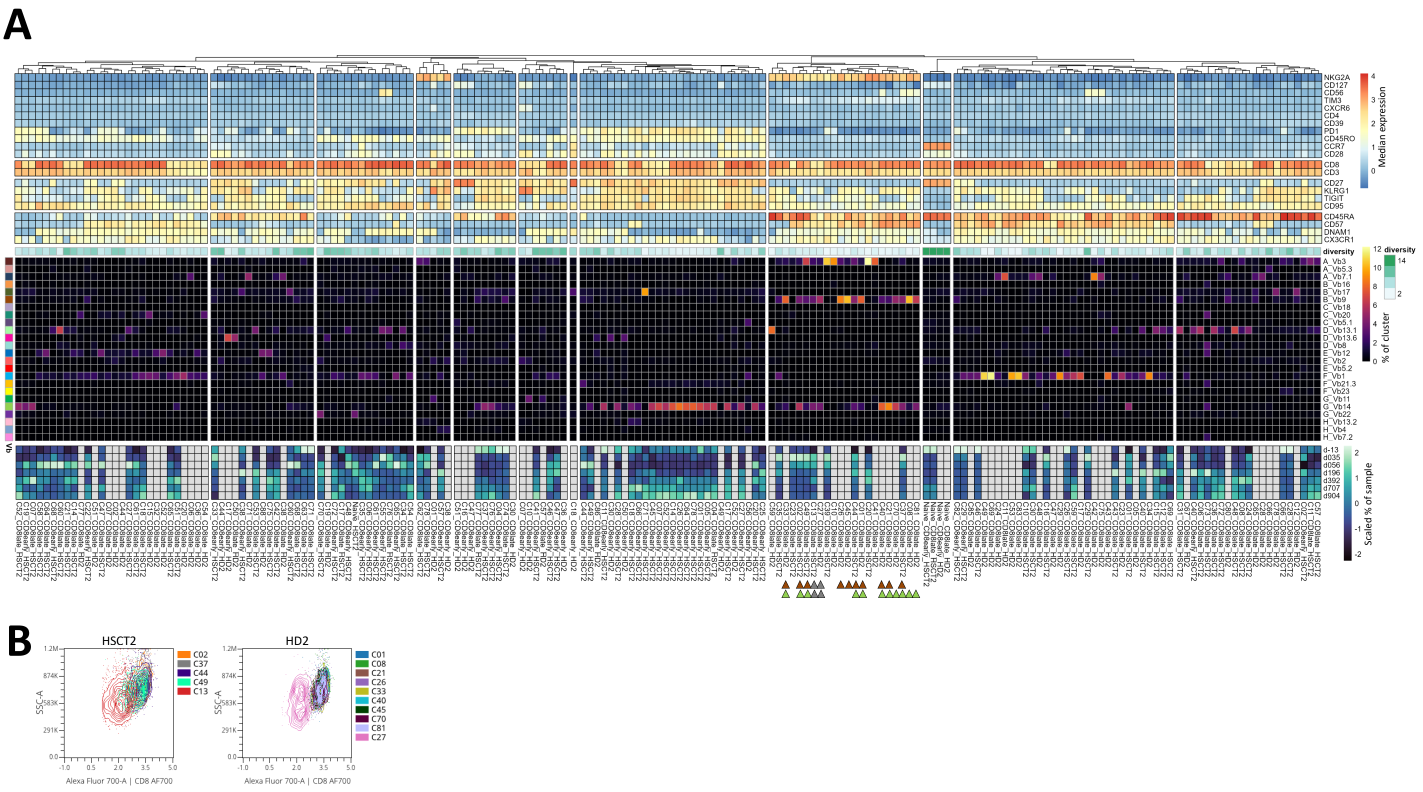


**Figure S11. Phenotype and frequency of Vβ dominant CD8 clusters of both patient HSCT2 and its donor HD2**

**A.** The CD8 early and late memory clusters that were defined as oligoclonal expansions in Figure 5 (HD2), 6 (HD2) and 8 (HSCT2) are included in the heatmap. The top heatmap represents the phenotype (median intensity), the second the TCR Vβ diversity (inverse Simpson index), the third the Vβ frequency and the bottom the scaled cluster frequency for the individual HSCT2 samples. As a reference the naïve CD8 T cells are included. The arrowheads point to the clusters as further discussed in Figure 8J-M. The color represents the TCR Vβ, except for the grey arrowheads which point to clusters removed from further analysis. **B.** Cluster C13-HSCT2 and C27-HD2 (grey arrowheads in A) were removed from further analysis for the TCR Vβ9 and Vβ14 quantification, since the low scatter and low CD8 indicate dying cells.
