## Supplementary table for "Immunophenotyping of T cells Combined with Vβ antibodies identifies long lasting CMV related T cell Expansions with a consistent TIGIT and PD-1 phenotype"

**Table S1 Donor and patient characteristics**

| | Sample | age | CMV status | CMV load plasma (log) | EBV status | Day post HSCT |
|---|---|---|---|---|---|---|
| **HC1** | 1 | 2 | + | NA | unknown | - |
| **HC2** | 1 | 17 | + | NA | + | - |
| **HC3** | 1 | 47 | + | NA | + | - |
| **HD1** | 1 | 38 | + | NA | + | - |
| **HD2** | 1 | 4 | + | NA | + | - |
| **HSCT recpient 1** | 1 | 15 | + | NA | + | -13 |
| **HSCT recpient 1** | 2 | 15 | + | 3.5 | + | 34 |
| **HSCT recpient 1** | 3 | 15 | + | 0 | + | 47 |
| **HSCT recpient 1** | 4 | 15 | + | NA | + | 204 |
| **HSCT recpient 1** | 5 | 16 | + | NA | + | 344 |
| **HSCT recpient 1** | 6 | 16 | + | NA | + | 666 |
| **HSCT recpient 2** | 1 | 1 | + | NA | + | -13 |
| **HSCT recpient 2** | 2 | 1 | + | 0 | + | 35 |
| **HSCT recpient 2** | 3 | 1 | + | 0 | + | 56 |
| **HSCT recpient 2** | 4 | 1 | + | NA | + | 196 |
| **HSCT recpient 2** | 5 | 2 | + | NA | + | 392 |
| **HSCT recpient 2** | 6 | 3 | + | NA | + | 707 |
| **HSCT recpient 2** | 7 | 3 | + | NA | + | 904 |

**Table S2 Antibodies**

**T cell panel spectral cytometry**

| | Specificty | | Antibody characteristics | | | | | |
|---|---|---|---|---|---|---|---|---|
| **Tube** | **CD designation** | **Alternative name** | **Fluorochrome** | **Isotype** | **Clone** | **Company** | **Catalog#** | **Dilution** |
| A | | Vb5.3 | PE | mouse IgG1 | 3D11 | Beckman Coulter | IM3497 | 10 |
| A | | Vb7.1 | PE+FITC | mouse IgG2a | ZOE | Beckman Coulter | IM3497 | 10 |
| A | | Vb3 | FITC | mouse IgM | CH92 | Beckman Coulter | IM3497 | 10 |
| B | | Vb9 | PE | mouse IgG2a | FIN9 | Beckman Coulter | IM3497 | 10 |
| B | | Vb17 | PE+FITC | mouse IgG1 | E17.5F3 | Beckman Coulter | IM3497 | 10 |
| B | | Vb16 | FITC | mouse IgG1 | TAMAYA1.2 | Beckman Coulter | IM3497 | 10 |
| C | | Vb18 | PE | mouse IgG1 | BA62.6 | Beckman Coulter | IM3497 | 10 |
| C | | Vb5.1 | PE+FITC | mouse IgG2a | IMMU157 | Beckman Coulter | IM3497 | 10 |
| C | | Vb20 | FITC | mouse IgG | ELL1.4 | Beckman Coulter | IM3497 | 10 |
| D | | Vb13.1 | PE | mouse IgG2b | IMMU222 | Beckman Coulter | IM3497 | 10 |
| D | | Vb13.6 | PE+FITC | mouse IgG1 | JU74.3 | Beckman Coulter | IM3497 | 10 |
| D | | Vb8 | FITC | mouse IgG2a | 56C5.2 | Beckman Coulter | IM3497 | 10 |
| E | | Vb5.2 | PE | mouse IgG1 | 36213 | Beckman Coulter | IM3497 | 10 |
| E | | Vb20 | PE+FITC | mouse IgG1 | MPB2D5 | Beckman Coulter | IM3497 | 10 |
| E | | Vb12 | FITC | mouse IgG2a | VER2.32 | Beckman Coulter | IM3497 | 10 |
| F | | Vb23 | PE | mouse IgG1 | AF23 | Beckman Coulter | IM3497 | 10 |
| F | | Vb1 | PE+FITC | rat IgG1 | BL37.2 | Beckman Coulter | IM3497 | 10 |
| F | | Vb21.3 | FITC | mouse IgG2a | IG125 | Beckman Coulter | IM3497 | 10 |
| G | | Vb11 | PE | mouse IgG2a | C21 | Beckman Coulter | IM3497 | 10 |
| G | | Vb22 | PE+FITC | mouse IgG1 | IMMU546 | Beckman Coulter | IM3497 | 10 |
| G | | Vb14 | FITC | mouse IgG1 | CAS1.1.3 | Beckman Coulter | IM3497 | 10 |
| H | | Vb13.2 | PE | mouse IgG1 | H132 | Beckman Coulter | IM3497 | 10 |
| H | | Vb4 | PE+FITC | rat IgM | WJF24 | Beckman Coulter | IM3497 | 10 |
| H | | Vb7.2 | FITC | mouse IgG2a | ZIZOU4 | Beckman Coulter | IM3497 | 10 |
| I | CD366 | TIM-3 | PE | rat IgG2a | 344823 | R&D | FAB2365P | 40 |
| I | CD45RO | CD45RO | FITC | mouse IgG1 | UCHL1 | DAKO | F0800 | 15 |
| A-I | CD186 | CXCR6 | BV421 | mouse IgG2a | K041E5 | Biolegend | 356014 | 40 |
| A-I | CD127 | IL7R | SB436 | mouse IgG1 | eBioRDR5 | Thermofisher | 62-1278-42 | 20 |
| A-I | CD57 | B3GAT1 | PB | mouse IgM | HNK1 | Biolegend | 359608 | 640 |
| A-I | CD3 | | BV480 | mouse IgG1 | UCHT1 | BD | 566105 | 640 |
| A-I | | TCRgd | BV510 | mouse IgG1 | 11F2 | BD | 745026 | 80 |
| A-I | CD45RA | | BV570 | mouse IgG2b | HI100 | Biolegend | 304132 | 160 |
| A-I | CD27 | TNFRSF7 | BV605 | mouse IgG1 | M-T271 | BD | 740398 | 40 |
| A-I | CD56 | NCAM1 | BV650 | mouse IgG2b | NCAM16.2 | BD | 564057 | 160 |
| A-I | CD279 | PD1 | BV711 | mouse IgG1 | EH12.1 | BD | 564017 | 40 |
| A-I | CD28 | | BV750 | mouse IgG1 | CD28.2 | BD | 747329 | 20 |
| A-I | | CX3CR1 | BV785 | mouse IgG2b | 2A9-1 | BD | 744489 | 80 |
| A-I | CD16 | FCGR3A | Vio515 | mouse IgG1 | REA423 | Miltenyi | 130-119-616 | 640 |
| A-I | CD4 | | AF532 | mouse IgG1 | SK3 | Thermofisher | 58-0047-42 | 40 |
| A-I | CD95 | FAS | PE-Dazzle594 | mouse IgG1 | DX2 | Biolegend | 305634 | 40 |
| A-I | CD226 | DNAM1 | BB700 | mouse IgG1 | DX11 | BD | 745864 | 40 |
| A-I | | KLRG1 | PerCP-ef710 | mouse IgG2a | 13F12F2 | Thermofisher | 46-9488-42 | 40 |
| A-I | CD39 | NTPDase1 | PE-Cy7 | mouse IgG1 | A1 | Biolegend | 328212 | 40 |
| A-I | CD159a | NKG2A | APC | mouse IgG2b | Z199 | BC | A60797 | 40 |
| A-I | | TIGIT | AF647 | mouse IgG1 | MBSA43 | eBioscience | 51-9500-42 | 80 |
| A-I | CD8 | | AF700 | mouse IgG1 | SK1 | Biolegend | 344724 | 160 |
| A-I | CD197 | CCR7 | APC-Fire750 | mouse IgG2a | G043H7 | Biolegend | 353246 | 40 |
